# *MGA* deletion leads to Richter’s transformation via modulation of mitochondrial OXPHOS

**DOI:** 10.1101/2023.02.07.527502

**Authors:** Prajish Iyer, Bo Zhang, Tingting Liu, Meiling Jin, Kevyn Hart, Jibin Zhang, Joo Song, Wing C. Chan, Tanya Siddiqi, Steven T. Rosen, Alexey Danilov, Lili Wang

## Abstract

Richter’s transformation (RT) is a progression of chronic lymphocytic leukemia (CLL) to aggressive lymphoma. *MGA* (*Max gene associated*), a functional MYC suppressor, is mutated at 3% in CLL and 36% in RT. However, genetic models and molecular mechanisms of *MGA* deletion driving CLL to RT remain elusive. We established a novel RT mouse model by knockout of *Mga* in the *Sf3b1*/*Mdr* CLL model via CRISPR-Cas9 to determine the role of *Mga* in RT. Murine RT cells exhibit mitochondrial aberrations with elevated oxidative phosphorylation (OXPHOS). We identified *Nme1* (Nucleoside diphosphate kinase) as a *Mga* target through RNA sequencing and functional characterization, which drives RT by modulating OXPHOS. As *NME1* is also a known MYC target without targetable compounds, we found that concurrent inhibition of MYC and ETC complex II significantly prolongs the survival of RT mice *in vivo*. Our results suggest that *Mga-Nme1* axis drives murine CLL-to-RT transition via modulating OXPHOS, highlighting a novel therapeutic avenue for RT.

**Statement of Significance:** We established a murine RT model through knockout of *Mga* in an existing CLL model based on co-expression of *Sf3b1*-K700E and *del*(*13q*). We determined that the *MGA/NME1* regulatory axis is essential to the CLL-to-RT transition via modulation of mitochondrial OXPHOS, highlighting this pathway as a novel target for RT treatment.

## Introduction

CLL is the most prevalent adult hematologic malignancy in North America(*1*, *2*). Richter’s transformation (RT), which occurs in up to 10-15% of patients with CLL(*3*), is a progression of the indolent CLL to an aggressive form of lymphoma, predominantly diffuse large B cell lymphoma (DLBCL). Patients who develop RT have a dismal prognosis, and median overall survival is less than 1 year, even with current moderate or high-intensity chemoimmunotherapy treatments(*4*). Moreover, RT patients following targeted therapy such as ibrutinib have a survival of about 4 months(*5*). Thus, understanding RT biology and designing better therapeutic approaches are urgently needed to treat this deadly disease.

Despite this fundamental clinical unmet need, a few obstacles impede the progress of RT study. First, RT diagnosis relies on clinical symptoms and pathological morphology changes in transformed cells. Lack of clear-cut definition, the rarity of RT samples, and the notion of similarity with aggressive CLL make identifying RT challenging(*6*). As a result, even though large-scale sequencing data is available in *CLL*(*7*, *8*), molecular mechanisms underlying CLL-to-RT transition are still limited. Second, the lack of human RT cell lines and limited animal models that faithfully recapitulate the human CLL-to-RT transition hamper mechanistic studies of RT and the testing of novel agents *in vivo*. Hence, interrogating pathways underlying this transition using novel murine models will facilitate our understanding of RT biology and uncover novel therapeutic targets.

A recent genetic characterization study using matched CLL and RT samples derived from 19 patients revealed that genetic lesions involved in cell cycle (89%, *TP53*, *CDKN2A/B*), MYC activation (74%, *MYC, MGA*), NOTCH pathway (32%, *NOTCH1, SPEN*), NF-kB signaling (74%, *BIRC3, EGR2*) chromatin remodeling (79%, *SETD2, SETD1A/B, ARID1A/B*) are enriched during CLL-to-RT transformation(*9*). The functional roles of *TP53/CDKN2A/B* deletions and *NOTCH* activation have been demonstrated to promote CLL to transform into aggressive lymphomas with DLBCL morphology(*10*, *11*). However, the roles of other genetic lesions during CLL-to-RT remains uncharacterized. In particular, loss-of-function mutations or deletions in *MGA (Max gene associated*), a MYC transcriptional repressor, are found in CLL at 3% but at 36% in RT, indicating a critical yet unknown role during CLL-to-RT (*7*, *9*, *12*). Here, we start with a murine CLL model based on the two most common genetic alterations found in human CLL samples, *Sf3b1* mutation and *Mdr* deletion [mimics del(13q)], and determine the impact of *MGA* deletion in the pathogenesis of RT. Our results highlight a novel *MGA*-driven regulatory axis driving CLL-to-RT transition and emphasize targeting this pathway as an effective treatment.

## Results

### *Mga* deletion leads to rapid CLL-to-RT transition upon engraftment into recipient mice

To determine the functional role of *Mga* in CLL and RT, we used a novel method to introduce genetic lesions into murine B cells, in which murine hematopoietic stem cells (Lin^-^cKit^+^Sca1^+^; hereafter LSK) were genetically edited *in vitro* and then adoptively transferred into sub-lethally irradiated CD45.1 mice. We first crossed a murine CLL line (*CD19*^*cre*/+^*Mdr*^*fl*/+^*Sf3b1* K700E^fl/+^)(*13*) with a mouse strain that conditionally expresses *Cas9*(*14*) to obtain a donor mouse line (*Cd19-Cre*^*fl*/+^ *Sf3b1*^*fl*/+^ *Mdr* ^*fl*/+^ *Cas9*^*fl*/+^). We isolated LSK cells (CD45.2^+^) from these mice and transduced them with lentivirus expressing non-targeting single guide RNA (sgRNA) or sgRNA targeting *Mga in vitro*. These *in vitro* edited LSK cells were then engrafted into sub-lethally irradiated recipient mice (CD45.1, n=15 per group) by tail-vein injection. CLL onset (B220^+^CD5^+^ CLL-like cells) was monitored by flow-cytometric analysis of peripheral bleeds bimonthly, starting at age 6 months and ending at 24 months (Fig. 1A). All the engrafted LSK cells achieved high successful engraftment along with high efficiency of *Mga* KO (Supplementary Fig. S1A-B). Our disease monitoring (14, 17 and 19 months) revealed a mixture of small and large cells based on forward and side scatter using peripheral blood-based flow cytometry (Fig. 1B, Supplementary Fig. S1C-D). B220 and CD5 were expressed by both small and large cells. Three out of 15 mice carrying *Mga* deletion showed clonal expansion of B220^+^CD5^+^ cells and leukemic infiltration of tissues by 21 months, while no mice in the control group developed CLL-like disease (Fig. 1B, Supplementary Fig. S1C-E). The observation of large cells in the peripheral bleeds led us to consider if the large B220^+^CD5^+^ cells are RT cells.

**Figure 1.**
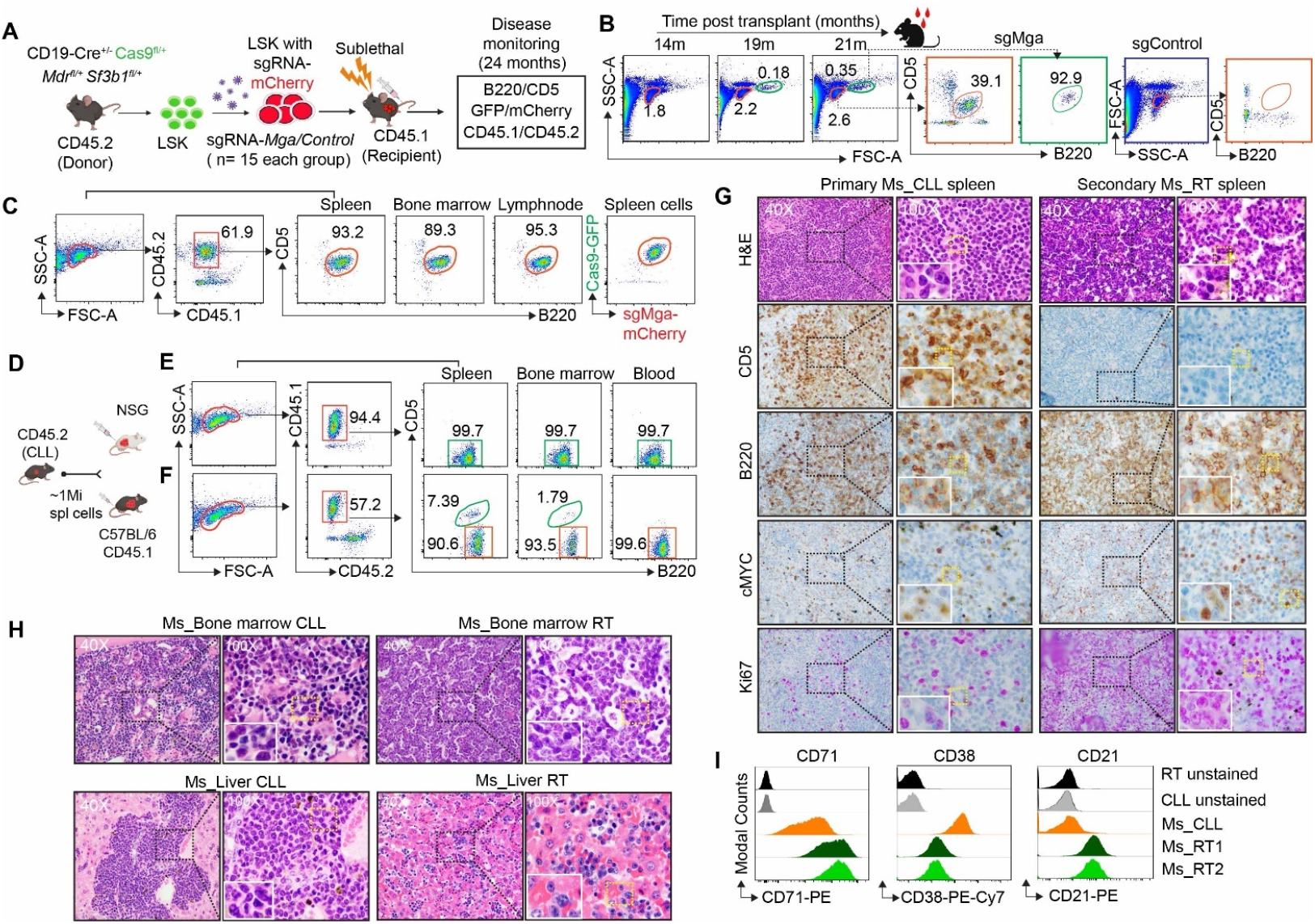
Knockout of *Mga* leads to Richter’s transformation in mice. **A,** Schema for generation of CLL-to-RT murine model. Genetically manipulated LSK (lin^−^c-kit^+^Sca-1^+^) hematopoietic progenitor cells derived from *Cd19-Cre* ^*fl*/+^*Cas9* ^*fl*/+^*Sf3b1*^*fl*/+^*Mdr* ^*fl*/+^mice (CD45.2) were transduced with lentivirus expressing sgRNAs against *Mga* or control were engrafted into sub-lethally irradiated CD45.1 recipient mice. The disease onset was monitored by examining the presence of B220^+^CD5^+^ cells from peripheral blood by flow cytometer. Transduced cells were identified based on mCherry expression. **B**, CLL disease in sg*Mga* group monitoring in the peripheral blood 14, 17, 19 months post-transplant by flow cytometry. Orange indicates the small lymphocytes and green indicates the large sized lymphocytes. Accumulation of B220^+^CD5^+^ (%) CLL-like cells is shown in the small and large lymphocytes. sgControl is also shown (Blue box). sgControl No CLL is shown in Orange box. **C,** Deletion of *Mga* leads to CLL. Accumulation of B220^+^CD5^+^(%) CLL-like cells in the spleen, bone marrow, and lymph node from the primary engrafted mice (CD45.2) was detected by flow cytometer 21 months post engraftment. **D,** Schema of engraftment of splenic B220^+^CD5^+^ CLL-like cells (derived from CD45.2) into either NSG or sub-lethally irradiated CD45.1 mice by tail-vein injection. **E-F,** Example of accumulation of B220^+^(%) cells in the spleen, bone marrow, and blood from the secondary NSG and CD45.1 mice. **G**, Immunohistochemistry staining of CD5, B220, MYC and Ki67 with H&E staining, on spleen sections from the primary (CLL) and secondary (RT) mice. 40X and 100X magnification images are shown. **H,** Representative H&E staining images from mouse bone marrow and liver in CLL and RT mice. 40X and 100X magnification images are shown. **I,** Flow cytometric analysis of CD71, CD38 and CD21 expression in two mouse RT cases and one CLL spleen case are shown.

We transplanted total splenic cells (Fig. 1C, mainly B220^+^CD5^+^) from the primary engrafted mice into either immunocompromised NSG or immunocompetent sub-lethally irradiated CD45.1 mice (secondary mice) (Fig. 1D). Transplantation of the total splenic cells resulted in a rapid expansion of B220^+^ B cells along with CD5 loss at 3 weeks post engraftment and leukemic infiltration in different lymphoid tissues (spleen, bone marrow, and lymph node) (Fig. 1E-F). All the mice had splenomegaly and infiltration in bone marrow as well as liver (Fig. 1E-H). Moreover, H&E staining of spleen section confirmed the presence of CLL-like cells with morphology highly similar to leukemia in the primary engrafted mice; however, upon secondary engraftment, these cells became larger with a morphology similar to aggressive lymphoma cells (Fig. 1G). Immunohistochemical staining with CD5, B220, Ki67, and MYC confirmed CLL identity in the primary mice and supported the presence of hyperproliferative and aggressive lymphoma in the secondary mice. We observed increased Ki67 and MYC in the aggressive secondary mice splenic B cells (Supplementary Fig. 2A-B). Of note, murine RT cells showed significant higher Ki67 proliferation index (>20%) than that of CLL cells (5-10%), indicating high rate of proliferation of RT cells. Flow cytometric analysis of murine RT cells revealed an increased level of CD71 and CD21 expression along with a decreased level of CD38 expression, consistent with previously reported RT cell characteristics(*15*, *16*) (Fig. 1I). The CLL-like and aggressive lymphoma-like cells were clonal based on immunoglobulin k (Igk) expression and shared the same immunoglobulin heavy chain gene variable region (*IGHV*) segment usage (Supplementary table 1). These observations indicated that *Mga* deletion coupled with *Mdr* deletion and *Sf3b1* mutation results in CLL transformation to RT, establishing a murine CLL-to-RT transition murine model.

### *Mga* deletion leads to activation of *MYC* and *NME1* in CLL and RT cells

To dissect the molecular mechanism underlying the CLL-to-RT transition, we performed RNA sequencing (RNA-seq) using RNA derived from splenic B cells from 4 mice without CLL, 2 mice with CLL, and their subsequent transformed RT cells from the secondary engraftment. Through differential gene expression analysis, we identified 424 CLL- and 1508 RT-associated upregulated genes, as well as 200 CLL-to-RT-associated upregulated genes (Fig. 2A). CLL dysregulated genes were highly enriched for MYC pathway targets and DNA repair while RT dysregulated genes were mainly enriched for MYC targets and inflammation pathways (Fig. 2B). Of note, CLL-to-RT transition was coupled with significant enrichment of MYC pathway targets and oxidative phosphorylation (OXPHOS) (Fig. 2C-D). When overlapping the upregulated (log2fc >1.5, p<0.05) gene list with murine CLL and RT datasets, we found 75 upregulated genes shared between CLL and RT groups (Fig. 2E), in which 3 genes (*MYBBP1A, NME1*, and *RPL22*) appear to be MYC targets directly regulated by *MGA* based on MGA binding protein and MYC targets overlap analysis(*17*). Among the three genes, *NME1* has been previously identified as transcriptionally regulated by *MGA* via promoter binding in lung adenocarcinoma(*17*). We confirmed the upregulation of *NME1* and *MYC* at the mRNA and protein levels in murine RT and CLL cells (Fig. 2F, Supplementary Fig 3A, 3C). We also validated *NME1* upregulation (~5-20 fold) in human CLL samples by RT-PCR and immunoblot (Fig 2G, Supplementary Fig. 3B). Notably, high *NME1* expression was significantly associated with inferior survival in patients with CLL and diffuse large B cell lymphoma (DLBCL) (Fig. 2H-I), indicating this gene’s critical yet unknown role in lymphomagenesis.

**Figure 2.**
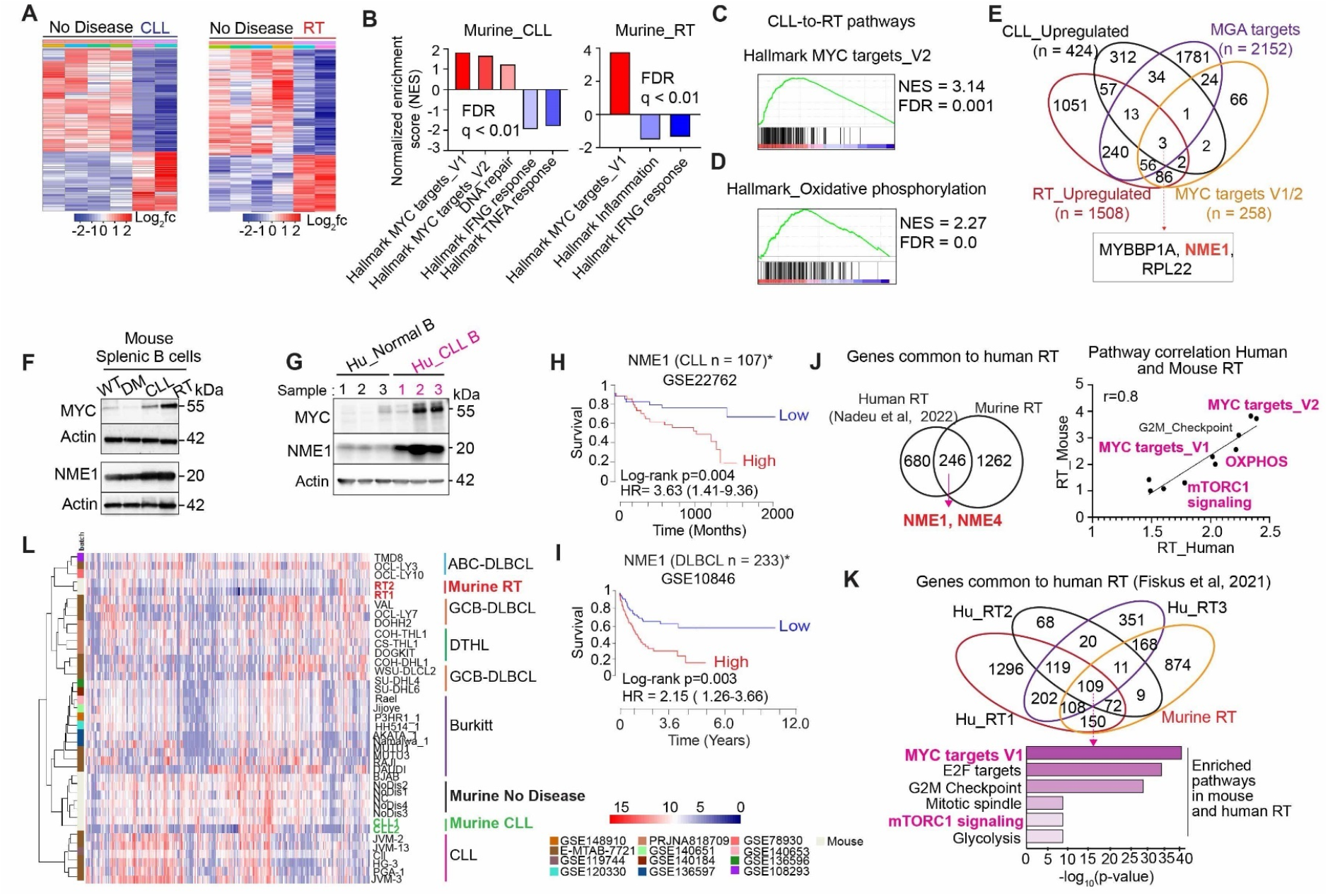
CLL-to-RT transition is coupled with *MYC* activation and increased *NME1* expression. **A,** RNA-seq was performed using RNA derived from splenic B cells either from mice without or with CLL (*Cd19-Cre*^+/−^*Sf3b1* ^fl/+^*Mdr*^fl/+^ *Cas9* ^fl/+^ *Mga*^-/-^), as well as mice transformed to RT (*Cd19-Cre*^+/−^*Sf3b1* ^fl/+^*Mdr*^fl/+^ *Cas9* ^fl/+^ *Mga*^-/-^). Heatmap illustrates differential gene expression among splenic B cells without CLL (n=4), with CLL (n=2), and with matched RT (n=2). Significantly upregulated and downregulated genes are shown in red and blue, respectively. **B-C,** Gene-set enrichment analysis (GSEA) of differentially expressed genes in murine CLL and RT identifies top dysregulated pathways ranked by normalized enrichment score (NES). **D,** Gene-set enrichment analysis of ranked genes from differential gene expression analysis between murine CLL and RT. **E,** Venn diagram shows the overlap of significantly dysregulated genes associated with CLL and RT with publicly available datasets, including Hallmark MYC targets V1/2 and *MGA* ChIP-Seq data A549 (ChIP-Atlas). *NME1* is one of the top *MYC* targets. **F,** Expression of MYC, NME1, and GAPDH were probed by immunoblotting using splenic B cells derived from wild type (WT), DM: No disease double mutant (*Cd19*Cre^fl/+^*Mdr* ^*fl*/+^ *Sf3b1* ^*fl*/+^ *Mga*^-/-^) Double mutant CLL and RT mice. **G,** Expression of MYC, NME1 and Actin were probed by immunoblotting using human normal B and CLL B cells. **H-I,** Kaplan-Meier survival curves for patients with CLL (**H**) or DLBCL(**I**) with high or low *NME1* expression. CLL (GSE22762) and DLBCL (GSE10846) datasets are from the *PRECOG (Prediction of clinical outcomes from Genomics) database. **J,** Venn diagram shows the overlap of murine-RT dysregulated genes with human CLL-RT dysregulated genes (Nadeu et al, 2022). Scatter plot showing the correlation of common dysregulated pathways in murine and human RT. **K,** Venn diagram shows the overlap of murine-RT dysregulated genes with human RT dysregulated genes (Fiskus et al, 2021). Pathway enrichment analysis from the common genes (n =109) between murine and human RT (n= 3). **L**, Heatmap representing hierarchical clustering of murine CLL-RT with human CLL and DLBCL cell lines.

To determine whether murine RT cells have a similar gene expression pattern to human RT cells, we overlapped murine RT-associated genes with three human RT cell datasets that were recently reported (*9*, *18*, *19*). From all three datasets, overlapped RT-associated genes revealed not only the upregulated genes but also the upregulation of pathways, including MYC, enrichment of MYC pathway targets, cell cycle, mTORC1 signaling, and glycolysis, indicating our murine RT model resembles human RT (Fig 2J-K, Supplementary Fig. 3D). In particular, upon overlapping our murine RT RNA-seq with a recently published human RT RNA-seq dataset (n = 19), we found *NME1* and *NME4* all upregulated in the human RT along with enrichment of MYC, OXPHOS, and mTORC1 pathways (Fig. 2J), indicating our murine RT cells shared similar gene expression pattern as human RT cells. Similarly, on overlapping the murine RNA seq dataset with another human RNAseq dataset from 3 samples, MYC targets V1 and mTORC1 were among the significantly enriched pathways (Fig. 2K). It is known that about ~90% of human RT cells are DLBCLs ABC type(*20*, *21*). To determine what type of human lymphoma our murine RT cells resembled, we compared gene expression of murine RT cells to 12 different types of human hematologic malignancy cell lines, including MYC-driven DLBCL, double-hit or triple-hit lymphoma (DTHL), and Burkitt’s lymphoma. The hierarchical clustering of the top 370 highly variable genes indicated that murine RT cells were similar to human ABC DLBCLs (Fig. 2L). In addition, murine CLL cells were also found to be closer to the human CLL cells, with gene expression signature distinct from non-tumor cells (Fig. 2L). Taken together, our findings show that murine RT cells resemble human RT cells with DLBCL gene expression signature, highlighting our murine RT model is faithful to human disease.

### RT cells have aberrant mitochondrial structural changes, increased oxidative phosphorylation, and metabolic reprogramming

Our gene expression analysis revealed MYC, OXPHOS and glycolysis pathways are enriched in RT cells and MYC is known to promote cell growth by regulating oxidative phosphorylation (OXPHOS) (*22*, *23*). Given the large size, aggressive growth, and increased MYC expression of RT cells, we hypothesize that *MGA* deletion drives CLL-to-RT transition through mitochondrial deregulation. Mitochondria are dynamic organelles, and alteration of respiration is usually coupled with aberrant ultrastructure, mtDNA, and ROS(*24*). In particular, mitochondria undergo changes in shape under oxidative stress (21,22). To evaluate mitochondrial structural alterations in RT, we performed electron microscopy-based morphometric analysis on murine splenic B cells. A greater percentage of mitochondria in the CLL group had aberrant mitochondria structures, including broken cristae with reduced width (Fig. 3A-C, Supplementary Fig 4B-C). Mitochondria in RT cells also exhibited larger cristae area and wider cristae (Fig. 3A-C, Supplementary Fig. 4A), indicating altered OXPHOS capability. Consistent with the morphological changes, we also detected increased protein expression in OPA1 and DRP1, two essential proteins regulating mitochondria fusion and fission (*25*), in both mouse (CLL and RT) and human (CLL) cells (Fig 3D-E, Supplementary Fig. 4B-C).

**Figure 3.**
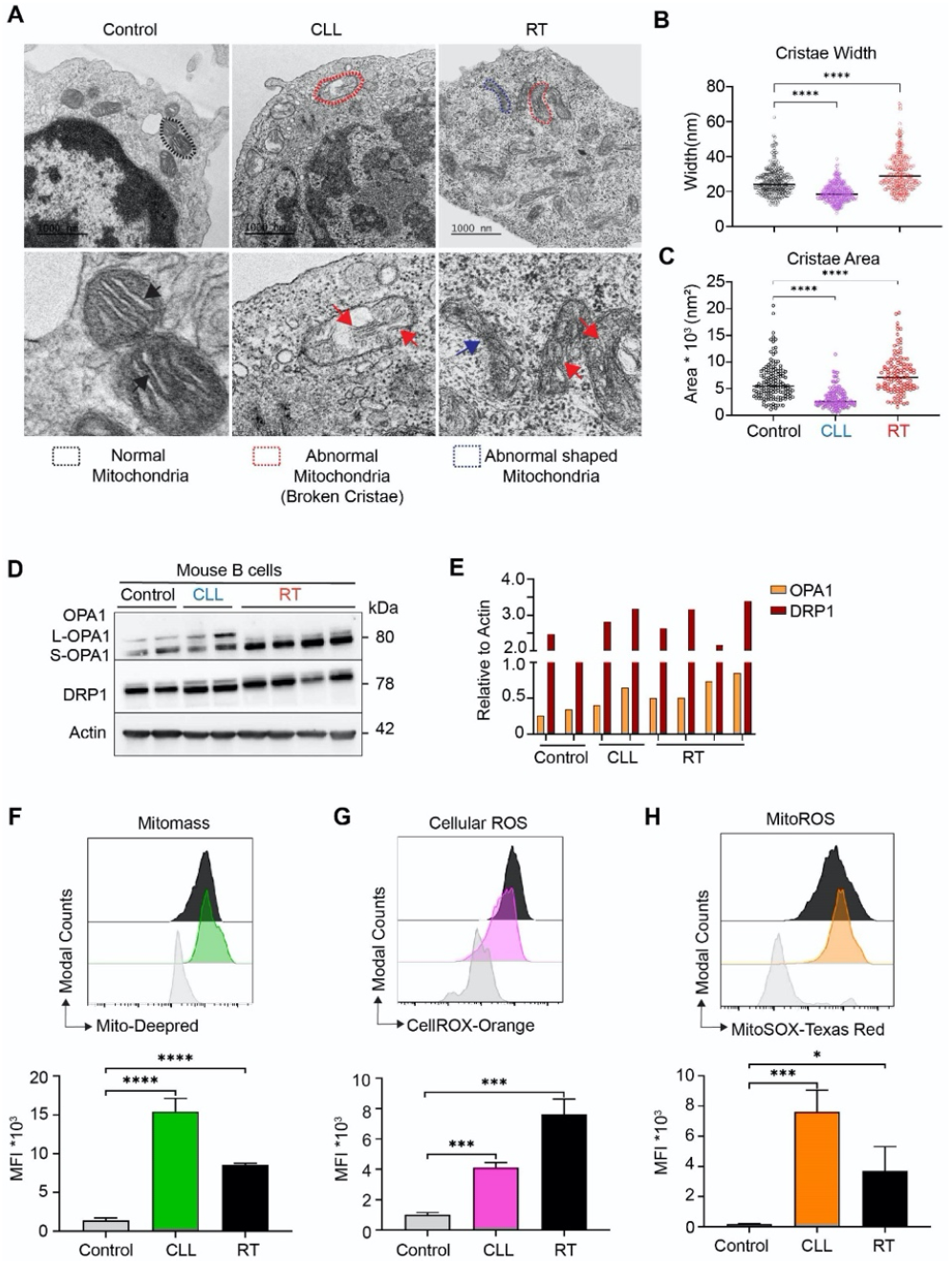
*Mga* deletion leads to increased mitochondrial dysregulation and abnormal cristae in murine CLL-RT. **A,** Representative electron micrographs of mouse splenic B cells from Control (No disease), CLL, and RT mice. (Scale bar-1000nm). Black and blue dotted lines indicate normal and abnormally shaped mitochondria. Black and red arrows show normal and abnormal cristae structures. **B-C,** Graph showing the quantification of cristae width (**B**) and cristae area (**C**) in murine control, CLL and RT. ****P < 0.0001 2-way ANOVA Dunnett’s multiple comparison test. **D,** Immunoblot showing the changes in mitochondrial fusion protein-OPA1 (L-OPA1 long isoform and S-OPA1 short isoforms) and fission protein-DRP1 in murine control, CLL and RT. **E,** Graph showing the quantification of OPA1, DRP1 in murine control, CLL and RT. **F-H,** Mitomass, cellular ROS, Mitchondrial ROS of splenic B cells derived from control (no disease), CLL, and RT mice were determined by MitoDeepred, CellROX Orange and MitoSOX based flow cytometry, respectively (n = 3 per group). MFI, mean fluorescence intensity. \*\*\*\**P*< 0.0001, \*\*\**P*<0.001, \**P*<0.05, 2-way ANOVA test.

The intact mitochondrial structure is essential to mitochondrial function and cell respiration. Given the distinct mitochondria structure changes, we further evaluated the mitochondrial mass and oxidative stress (cellular and mitochondrial ROS) in splenic B cells derived from control, CLL, and RT mice using flow cytometry-based assays. Murine CLL and RT cells showed higher mitochondrial mass as compared to control normal B cells. Similarly, mitochondrial ROS was higher in murine CLL and RT cells as compared to control normal B cells. Cellular ROS was higher in RT cells as compared to CLL and normal B cells. These data suggest that *Mga* deletion leads to metabolism reprogramming. Of note, both murine and human RT cells had upregulated GLUL expression (Glutamine ammonia ligase) (Supplementary Fig. 4D-E), a glutamine synthetase that is transcriptionally regulated by MYC and supports nucleotide synthesis, amino acid transport, and TCA cycle(*26*). Taken together, in addition to shared gene and protein expression with human RT cells, *Mga* deleted murine RT cells display dramatic mitochondria ultrastructure alteration, enhanced OXPHOS, and metabolic reprogramming.

### Knockout of *MGA* in human cell lines leads to abnormal cristae structure, hyperactive mitochondrial activity, and hyperproliferation, recapitulating features of murine RT cells

Given that our murine model strongly implicates mitochondrial function is altered in RT cells and may be involved in the onset of RT, we set out to investigate the molecular circuitry changes underlying *MGA* deletion-associated mitochondrial dysfunction in human B lymphoid cell lines. We knockout (KO) *MGA* in several human cell lines (CLL: MEC1, HG3; acute lymphoblastic leukemia: Nalm6E) by CRISPR/Cas9 technology to determine whether *MGA/MYC/NME1* axis can be recapitulated in human cells (Supplementary Fig. 5A). *MGA* KO Nalm6E and MEC1 cells exhibited higher oxygen consumption rate (OCR) at baseline and over time in response to modulators of the mitochondrial electron transport chain (ETC) and OXPHOS (Fig. 4A-B, Supplementary Fig 5A-B). *MGA* KO Nalm6E cells also showed dependency on all the three energy sources measured by Seahorse substrate oxidation test (glucose: UK5099 glutamine: BPTES lipids: Etomoxir (ETO)) (Supplementary Fig. 5C-D), indicating the substrates underlying higher OXPHOS. Of note, these cells showed consistently higher steady-state levels of cellular ROS (p < 0.001) coupled with increased mitochondrial DNA (mtDNA) copy number and amount of ATP (Fig. 4C-E) (p < 0.001). The mitochondrial membrane potential was also significantly elevated upon *MGA* KO (Supplementary Fig. 5E), consistent with the hyper-energetic state observed in murine RT cells. These observations support that *MGA* deletion impairs mitochondrial quality control processes. Likewise, we also examined the impact of *MGA* deletion on mitochondria function in other B cell lines we generated. Similar to our observation in Nalm6E cells, *MGA* deletion induced a consistent increase in cellular ROS production in MEC1 and HG3 cells (Supplementary Fig. 5F), indicating that mitochondrial dysregulation is a general feature induced by *MGA* deletion.

**Figure 4.**
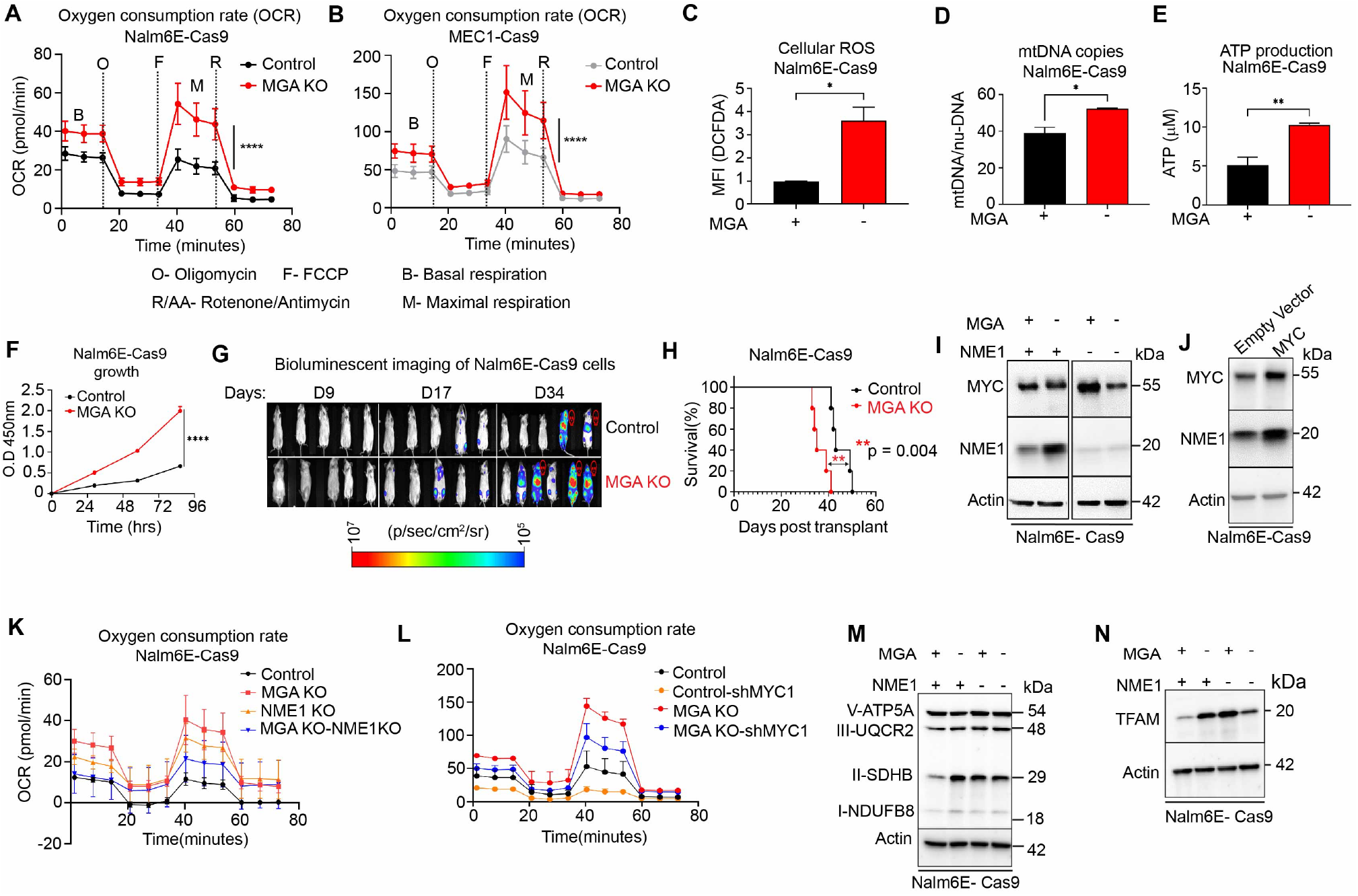
*MGA* deletion leads to increased OXPHOS and abnormal cristae structure. **A-B,** OCR was determined by Seahorse Mitostress analysis in Nalm6E-Cas9 cells, MEC1- Cas9 with and without *MGA*. **C-D,** Cellular ROS and mitochondria DNA (mtDNA) in Nalm6E cells with or without *MGA* were determined by H_2_DCFDA flow-based and PCR assays, respectively. **E,** Cellular ATP determined by Cell titer glo assay in Nalm6E cells with or without *MGA*. Y axis shows the ATP concentration (μM). **F**, In vitro growth monitoring for 4 days by CCK8 assay in Nalm6E-Cas9 cells with or without *MGA*. Y axis shows the absorbance at 450nm. \*\*\*\**p*< 0.0001, by 2-way ANOVA. **G**, *In vivo* bioluminescent imaging of NSG mice with engraftment of control (n = 5) and *MGA* KO (n=5) Nalm6E cells at day 9, 17, and 34 post engraftments. All the images were acquired at the same scale. **H,** The survival of NSG mice xenografted with control and *MGA* KO Nalm6E cells were measured and expressed as a percent of the initial number of mice in each group. Log-Rank test was used. **I,** Expression of MYC and NME1 in Nalm6E-Cas9 cells with different genetic lesions detected by immunoblotting. **J,** Expression of MYC, NME1 in Nalm6E-Cas9 cells overexpressing MYC detected by immunoblotting. **K-L,** Oxygen consumption rate measured in Nalm6E cells (Control and *MGA* KO) with *NME1* KO **(K)** or *MYC* knockdown **(L)** by Seahorse Mitostress assay. Y axis represents the oxygen consumption rate, and the X-axis represents the time of the experiment**. M**, ETC proteins in Nalm6E-Cas9 cells with different genetic lesions**. N,** Expression of TFAM (transcription factor A, mitochondrial) in Nalm6E-Cas9 cells with different genetic lesions detected by immunoblotting. \*\*\*\**p*< 0.0001, \*\*\**P*<0.001, \*\**P*<0.01, \**P*<0.05.

To determine the consequence of the *MGA* deletion-induced oxidative stress, we engrafted Nalm6E control, and*MGA* KO cells with stable overexpression of luciferase into immunodeficient NSG mice. Through real-time bioluminescent imaging monitoring over 34 days, we found that *MGA* KO accelerated cell proliferation *in vivo* (Fig. 4F). As a result, mice with engraftment of *MGA* KO cells had poor overall survival compared to control cells (Fig. 4G, p=0.00; log-rank test). Taken together, *MGA* deletion led to mitochondria structure change, higher levels of OXPHOS and ROS, ATP amount, and cell hyperproliferation, confirming a solid linkage between mitochondrial dysregulation and growth advantage induced by *MGA* deletion.

### *MGA* deletion modulates OXPHOS via MYC and NME1

To elucidate the molecular regulatory network underlying *MGA* deletion in driving mitochondrial dysregulation and cell growth, we generated *NME1* KO Nalm6E cells with and without *MGA* (Supplementary Fig. 5A). We examined the expression of *MYC* and *NME1* at both RNA and protein levels. Loss of *MGA* in Nalm6E increased *NME1* expression (Fig. 4H, Supplementary Fig. 5A), supporting *MGA* as a negative regulator for *NME1*. We explored the effect of *MGA* KO in other B cell lines and found NME1 was distinctly upregulated at the protein level in MEC1, validating *NME1* as a target of *MGA* (Supplementary Fig. 6C). We also discovered that MYC was upregulated in Nalm6E cells upon *NME1* deletion at both RNA and protein levels, suggesting *NME1* is a negative regulator for MYC (Fig. 3G, Supplementary Fig. 6A). We found no drastic change in overall MYC protein expression in Nalm6E cells upon *MGA* deletion (Fig. 3G, Supplementary Fig. 6B), possibly owing to the level of increase of NME1 expression upon *MGA* deletion that inhibits MYC expression (Supplementary Fig. 6B), highly suggestive of a tight regulation within this axis, however on further fractionation of cytosolic and nuclear isolates, we observed increase in nuclear MYC (Supplementary Fig 6B). While *MGA* deletion in MEC1 cells increased MYC, *NME1* KO failed to further increase MYC expression (Supplementary Fig. 6C), possibly due to cell line specific effects. We observed a subtle increase in MYC and NME1 in HG3 cells upon *MGA* deletion (Supplementary Fig. 6D). As increased MYC has been associated with proliferation related pathways such as pERK and mTORC1(*27*, *28*), we detected increased pERK1/2 and mTORC1 signaling (p4E-BP1) cells with *MGA* deletion in MEC1 and HG3 cells (Supplementary Fig. 6E). Increased mTORC1 signaling was not accompanied with increased AKT signaling (Supplementary Fig. 6F). Notably, deletion of *MYC* significantly impacted cell growth in all cells (Supplementary Fig. 6G), while *NME1* KO affected cell growth in *MGA* KO cells (Supplementary Fig. 5H).

To dissect the relationship between *MYC* and *NME* in *MGA* deletion-mediated mitochondria dysregulation, we performed Seahorse Mito stress assay on Nalm6E cells with or without *NME1* or *MYC. NME1* KO cells appeared to have an increased basal and maximal oxygen consumption rate compared to control cells, possibly owing to increased MYC in these cells (Fig. 4K). *MGA* KO cells showed a dependency on *NME1* for OXPHOS (Fig. 4K). Concurrent KO of *MGA* and *NME1* resulted in partial normalization of metabolic activity, consistent with our notion that *NME1* is required for cell proliferation in *MGA* KO cells (Supplementary Fig. 6H). In line with the role of *MYC* in promoting high OXPHOS in *NME1* KO cells, control cells demonstrated a drastic reduction for basal and maximal oxygen consumption rates upon MYC deletion, while *MGA* KO cells showed dependency on *MYC* for OXPHOS (Fig. 4L). These results reveal that *MGA* deletion shifts OXPHOS from an MYC-dependent mode to an MYC and NME1 co-dependent mode.

To determine the molecular basis for *MYC/NME1* OXPHOS dependency in *MGA* KO cells, we examined the OXPHOS mitochondrial ETC complex and TFAM (transcription factor A, mitochondrial), a key regulator for mitochondrial DNA replication(*29*). Mitochondrial ETC complex is the machine used for cells to generate ATP from the oxidation of carbohydrates, fats, and proteins and is composed of five separate protein complexes [I (NDUFB8), II (SDHB), III (UQCRC2), IV (MTCOX1), V (ATP5α)](*30*). *NME1* loss resulted in a substantial decrease in subunit I, while *MGA* KO cells increased subunit II (Fig. 4M). Increase in complex II in *MGA* KO Nalm6E and decrease in complex I in NME1 KO cells was further confirmed using immunoblotting on fractionated mitochondrial protein isolates (Supplementary Fig 6I). The increase of complex II was also seen in MEC1 cells with *MGA* KO (Supplementary Fig. 6J). Concurrent deletions of *MGA* and *NME1* led to a subunit II downregulation and a subunit I reduction compared to control cells, which could form the basis of *NME1* dependent OXPHOS in *MGA* KO cells (Fig. 4M). Moreover, we observed an *NME1*- and *MYC*-dependent TFAM expression pattern in *MGA* KO cells (Fig. 4N, Supplementary Fig. 6K). This distinct mitochondrial ETC complex and TFAM expression pattern upon *MYC* or *NME1* deletion corroborated our observation that both *NME1* and *MYC* have roles in regulating mitochondrial OXPHOS in *MGA* KO cells.

### AZ5576 and TTFA treatment provide therapeutic benefits for RT *in vivo*

The MYC and NME1 co-regulation mode of OXPHOS upon *MGA* deletion provides RT cells a maximum potential for metabolic reprograming, however, it also posits a challenge for treating RT disease. Thus, better treatment of RT is likely to be achieved through targeting mitochondrial dysregulation in *MGA* KO-induced disease. We, therefore, tested inhibitors of ETC complex II (TTFA: Thenoyltrifluoroacetone) and CDK9 (AZ5576) in the Nalm6E cell line (Fig. 5A-B). TTFA and CDK9 inhibited the expression of complex II/succinate dehydrogenase, MYC, NME1, pERK, and mTOR (both phospho and total 4E-BP1) pathways in Nalm6E and MEC1 cells (Fig. 4A-B, Supplementary Fig.7A-D), suggesting perturbation of mitochondrial dysregulation could impact cell proliferation.

**Figure 5.**
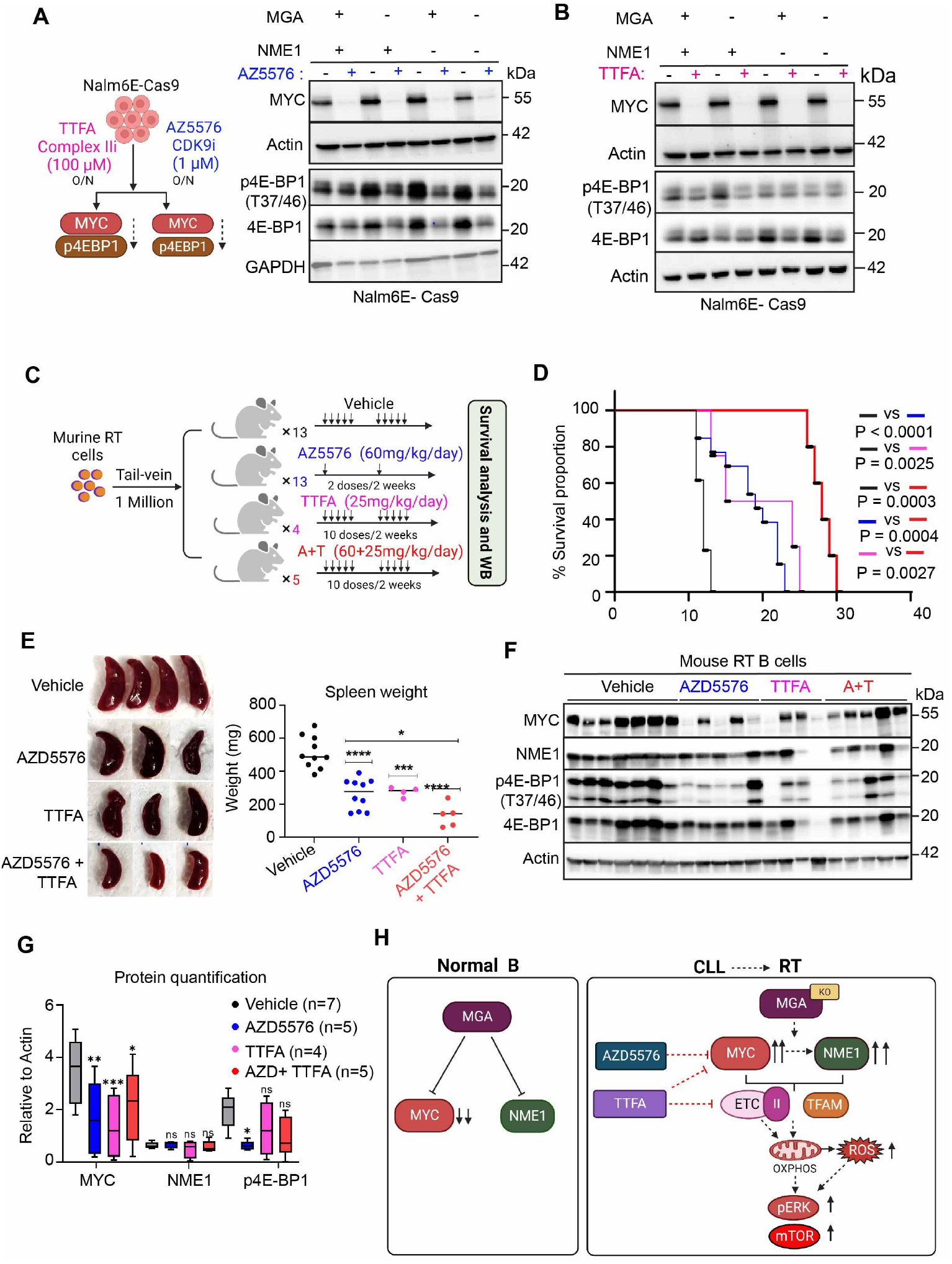
AZ5576 and TTFA prolong the survival of RT mice by inhibiting MYC and NME1 expression. **A-B,** MYC, p4E-BP1, and total 4E-BP1 expression detected by immunoblotting in Nalm6E-Cas9 cells treated with CDK9i AZ5576 (1μM) (A) or complex II TTFA (100 μM) (B) overnight. **C,** Schema for *in vivo* drug treatment experiments. **D,** Kaplan-Meier survival curves of RT mice treated with AZ5576 or TTFA alone or in combination. Log-Rank test. **E,** Images of spleens and spleen weight (mg) from RT mice with different treatments were collected at the endpoint. **F,** Immunoblotting of MYC, NME1, p4E-BP1, and 4E-BP1 protein in splenic B cells derived from RT mice at the endpoint. **G,** Protein quantification of MYC, NME1, and p4EBP1 in different groups with respect to loading control from panel F,\*\*\*\**P*< 0.0001, \*\*\**P*<0.001, \*\**P*<0.01, \**P*<0.05, ^ns^ Not significant. **H,** Model for the molecular mechanism of *MGA* KO driven CLL-to-RT transition.

We then explore whether these inhibitors can provide therapeutic benefits for RT mice *in vivo*. We first engrafted RT cells into NSG mice and established stable RT mice in 2 weeks (Supplementary Fig. 7E). Then we started the treatment with oral administration of either vehicle (n = 13); AZ5576 (60mg/kg) (n = 13); TTFA (25mg/kg) (n = 4); and AZ + TTFA (60 and 25mg/kg) (n = 5) into these mice. AZ5576 was administered once a week, while TTFA was given 5 days a week (Fig. 4C). AZ5576 and TTFA treatment significantly prolonged the survival of RT mice (p=0.0007) with a significant (p < 0.0001) reduction in total spleen weight. AZ+TTFA combination was further effective in increasing the survival of RT mice by 10 days (p=0.0003) and reducing spleen weight significantly (p<0.0001) as compared to single treatments. (Fig. 5D-E). AZ5576 and TTFA treated RT mice B cells had reduced MYC, NME1, SDHB, SDHA (complex II), NDUFB8 (complex I), ATP5A (complex V) at the RNA level (Supplementary Fig. 7F). RT cells derived from mice with AZ5576 and TTFA treatment had reduced MYC, NME1, and phospho-4E-BP1 T37/46 protein expression (mTOR pathway) (Fig. 5F-G), confirming the effective targeting of the *MGA/MYC/NME1* axis *in vivo*. Taken together, concurrent targeting of MYC and OXPHOS provides synergistic effects in treating RT *in vivo*, highlighting *MGA/MYC/NME1* regulatory axis as a novel target for RT treatment.

## Discussion

Here, we established a novel RT murine model faithful to human genetics through silencing *Mga* in an existing CLL model (Fig. 4H). Through extensive characterization of this murine RT model and functional experiments of human cell lines with *MGA* deletion, we discover that the *MGA-NME1* axis drives RT through mitochondrial OXPHOS upregulation, and concurrent targeting of MYC and OXPHOS pathways provides therapeutic benefits for RT mice *in vivo*, suggesting a new therapeutic avenue for RT patients.

Genetic manipulation recently established several RT models based on an aggressive Eμ-TCL1–transgenic CLL mouse model. These recent models include B-cell-specific deletion of *TP53/CDKN2A/CDKN2B*, deletion of *NFAT2* (downregulation upon deletions of *CDKN2A* or *TP53*), and overexpression of activated *AKT*, activated *NOTCH1*, or *c-MYC* to develop aggressive lymphomas with diffuse large B-cell lymphoma morphology (*10*, *11*). Instead of using Eμ-TCL1–transgenic mouse, an aggressive form of CLL mouse model, we started with an indolent CLL mouse model that fully recapitulates human CLL genetics (13q deletion and *SF3B1* mutation) as these two genetic lesions are the most common co-occurring events in CLL. Although CLL cells derived from these mice displayed increased MYC and NME1 protein expression, transformation into RT was barely observed by 24 months. However, CLL cells rapidly transformed into RT upon engraftment of *MGA*-deficient cells into recipient mice within 2 weeks, irrespective of host immune status, suggesting that cellular stress during the engraftment process is essential to the rapid transition from CLL to RT. Cross-validation of gene expression between murine RT, primary human RT cells, as well as human hematologic cell lines corroborated that our murine model faithfully recapitulates human RT features with DLBCL gene expression feature.

*Mga* deletion-induced mitochondrial ROS accumulation and metabolic reprogramming underlie CLL-to-RT transition. In CLL, mitochondria are dysregulated, with increased oxidative phosphorylation as the vital source of ROS(*31*). However, until now, the underlying mechanisms of oxidative stress in CLL and their contributions to RT onset are elusive. Our results support that *MGA/MYC/NME1* axis induces extensive mitochondria structure changes, increased OXPHOS, ROS accumulation by metabolic reprogramming through upregulation of ETC complex II and TFAM that eventually leads to activation of mTOR pathway and ERK pathway to contribute to cell proliferation. Consistently, a recent longitudinal study of CLL to RT samples (n = 54), identified a (OXPHOS)^high^ B cell receptor (BCR)^low^ transcriptional signature and showed OXPHOS as a potential therapeutic target in RT(*32*). Also, a recent multi-omics analysis of primary CLL samples suggests that mTOR-MYC-oxidative phosphorylation axis is significantly associated with an aggressive type of CLL(*33*). Our murine model and cell line characterization recapitulates the observation of MYC-OXPHOS-mTOR activation in human hyperproliferative CLL and further suggests that this axis underlies CLL-to-RT transformation.

Our results further suggest *NME1* has a critical role in metabolic reprogramming upon *MGA* deletion. We discover that *NME1* is a negative regulator of *MYC* and a direct target of *MYC* in cells without *MGA* deletion (Fig.4I-J). The co-dependent mode on *MYC* and *NME1* in *MGA* KO cells maximizes the cell’s potential to tolerate ROS accumulation and activate downstream oncogenic signaling, promoting cell growth via modulating mitochondria function. How NME1 interacts with mitochondria is not well-explored. NMEs directly interact with and channel GTP to specific dynamin-related GTPases to drive membrane remodeling and mitochondrial fusion (*34*, *35*). It has been reported that long-chain fatty acyl CoA (LCFA-CoA) inhibits NME1/2 (*36*). Our results also indicated that NME1 regulates a few mitochondrial genes, suggesting NME1 may not directly interact with mitochondria. Future molecular studies and biochemistry assays are anticipated to determine the mechanism of how *NME1* exerts its impact on mitochondria.

Targeting *MGA*/*MYC*/*NME1* axis provides therapeutic benefits for RT. Our novel murine RT model allowed us to explore new treatment options for this aggressive disease. Although inhibitors for either AZ5576 or TTFA can improve the survival of RT mice *in vivo*, combination treatment demonstrates a synergistic effect, suggesting cellular metabolic function is a tractable target in RT.

## Acknowledgments

This work was supported by grants from the National Institutes of Health (NCI) R01CA216273 and R01CA21623 (to LW), and R01CA244576 and the Leukemia and Lymphoma Society Translational Research Program Award 6517-22 (to AVD). AVD is a Leukemia and Lymphoma Society Scholar in clinical research. We also acknowledge Analytical Cytometry Core and Small Animal Imaging Core at City of Hope supported by the National Cancer Institute of the NIH under award number P30CA033572. We thank D. Lynne Smith, Ph.D., for carefully reading the manuscript and for editorial help.

## Methods

### Animals

All animals were housed at the City of Hope National Medical Centre (COH). All animal procedures were completed in accordance with the guidelines for the Care and Use of Laboratory Animals. All protocols were approved by the Institutional Animal Care and Use Committees at COH. To obtain heterozygous expression of *Sf3b1* mutations and *Mdr* deletion in B cells, we crossed *Sf3b1-K700E* floxed mice(*37*) with *Mdr* floxed mice(*38*) to generate *Sf3b1*^*fl*/+^*Mdr*^fl/fl^ mice, which were then crossed with CD19Cre (*Cd19*-Cre^+/+^) to obtain double mutant mice(*Cd19*-Cre^+/−^*Sf3b1* ^*fl*/+^*Mdr*^*fl*/+^). The double mutant mice were further crossed with CRISPR-Cas9 GFP mice (*14*) to obtain (*Cd19*-Cre^+/−^*Sf3b1* ^*fl*/+^*Mdr*^*fl*/+^ *Cas9* ^*fl*/+^).

### Murine splenic B cell isolation

Mice were euthanized in a CO_2_ chamber, and spleens were harvested. Spleens were dissociated to form a single-cell suspension, and red blood cells were lysed. B cells were immunomagnetically separated using the MACS mouse B cells isolation kit (Miltenyi Biotec, Auburn, CA) and L.S. columns (Miltenyi Biotec, Auburn, CA).

### Cell culture and reagents

Nalm6 (pre-B ALL cell line) and HG3 (CLL cell line), MEC1 (CLL cell line), were cultured in RPMI media supplemented with 10% FBS with 1% penicillin and streptomycin. All cell lines were incubated at 37°C with 5% CO_2_. These cell lines were lentivirally transduced with SpCas9 (Addgene: #52962, Watertown, MA) and selected with blasticidin (InvivoGen, San Diego, CA). The adherent cells HEK293T-LentiX (TAKARA, Mountain View, CA) were grown in DMEM (Sigma, St. Louis, MO) with 10% FBS and 1% penicillin and streptomycin. Nalm6 isogenic cell line *SF3B1*-K700E (Nalm6E) was purchased from Horizon discovery and previously reported (Darman et al., 2015). All cell lines were authenticated by STR analysis and determined as mycoplasma free before being used for experiments.

### Virus production and transduction

HEK293T-lentiX cells are used for lentivirus production. HEK293T-lentiX cells were plated on a 6-well plate at 3 × 10^5^ cells per well in DMEM medium with 10% FBS and allowed to adhere overnight. All transfections were carried out using polyethyleneimine (PEI Max 40K #24765, Polysciences, Warrington, PA) reagent at a 4:2:3 ratio of sgRNA/overexpression-construct: pVSVG: psPAX2 in Opti-MEM media (Life Technologies, Thermo Fisher Scientific Inc, Waltham, MA). Viral supernatants were collected 48 hours post-transfection and filtered through a 0.45 μm Nalgene syringe filter SFCA (Whatman, Clifton, NJ) before being concentrated by ultracentrifugation in 38.5mL tubes (#344058, Beckman Coulter, Brea, CA) at 22000 x g for 2 hours at 4°C. Cells (2×10^5^) were spin transduced with 10-20μl of the concentrated virus at 37°C at 2200*g* for 90 minutes with polybrene reagent (8-10μg/ml, Millipore Sigma, Billerica, MA). The cells were washed with PBS and resuspended in fresh RPMI media supplemented with 10% FBS with 1% penicillin and streptomycin 24 hours post-transduction. sgRNAs targeting either *MGA* or *NME1* was cloned in pLKO5.sgRNA.EFS.mCherry or pLKO5.sgRNA.EFS.GFP (Addgene, Watertown, MA), respectively. mCherry or GFP single or double-positive cells were sorted 10-15 days post-transduction, and stable cell lines were maintained with more than 95% positivity.

### Functional analysis of mitochondria

#### Measurement of Cellular ROS

To measure the levels of ROS in the cytoplasm, 1×10^5^ cells were incubated with H_2_-DCFDA (non-fluorescent dye, Abcam, Boston, MA)/ CellROX Orange (Thermo Fisher Scientific Inc, Grand Island, NY) in PBS (10μM) for 30 minutes at 37°C under 5%CO_2_. H_2_-DCFDA reacts with hydrogen peroxide and gives green fluorescence (DCFDA). CellROX Orange is detected at absorption/emission of 545/565nm. Cells were then washed with PBS and analyzed by flow cytometry.

#### Measurement of Mitochondrial membrane potential

To measure mitochondrial membrane potential, 1×10^5^ cells were incubated with membrane potential sensitive cyanine dye DilC_1_(5) (MitoProbe DilC_1_(5) assay kit, Thermo Fisher Scientific Inc, Grand Island, NY) in PBS (50nM) for 30 minutes at 37°C under 5%CO_2_. The dye primarily accumulates in mitochondria with active membrane potential. Cells were then washed with PBS and analyzed by flow cytometry. Cells were then washed with PBS and analyzed by flow cytometer (633nm-Far red).

##### Measurement of mitochondrial volume/mass

To measure mitochondrial mass, cells were first spin down at 1500g for 5min and resuspended in 100μl of PBS (Phosphate buffer saline) containing 0.2μM Mitotracker green (Dojindo Mol Tech)/ 0.1μM MitoBright LT DeepRed (Dojindo Mol Tech). Suspended cells were incubated for 30 mins under 5% CO_2_ incubator at 37° C. Cells were washed with PBS and analyzed by flow cytometry using a FITC (Fluorescein isothiocyanate) filter.

##### Measurement of mitochondria ROS

To measure the levels of ROS in mitochondria, we perform cell staining with MitoSOX (Thermo Fisher Scientific Inc, Grand Island, NY) in PBS for 30min at 37°C under 5%CO_2_. MitoSOX gives red fluorescence on reacting with mitochondria dependent ROS. Cells are washed with PBS and analyzed by flow cytometry.

#### Cellular respiration measurement

Cells were plated in a 6-well plate and incubated in RPMI with 10% FBS under 5% CO_2_ at 37°C 48 hours before the Seahorse experiment. On the day of assay, 8×10^4^ viable cells per well were seeded in poly-lysine-coated (Sigma-Aldrich, St. Louis, MO) XFe96 Seahorse Cell Culture Microplates (Agilent Technologies, Santa Clara, CA). Plates were centrifuged at 300g for 5 minutes with no brake and incubated for 30 minutes with no CO_2_ at 37°C. The following procedures were carried out with the Seahorse XFe96 Oxidative Stress Assay (103015-100, Agilent Technologies, Santa Clara, CA). First, basal cellular respiration was recorded by simultaneously measuring oxygen consumption rate (OCR) and extracellular acidification rate (ECAR). Next, 1.5 *μ*M oligomycin (ATP synthase inhibitor) was injected to stop ATP-linked respiration, leading to a direct measurement of proton-leak and non-mitochondrial oxygen consumption. After that, 1.0 μM FCCP was injected to induce maximal respiration/uncoupled oxidative phosphorylation (OXPHOS). Lastly, 0.5 μM rotenone and 0.5 μM antimycin A (Complex I and Complex III inhibitors, respectively) were injected, inhibiting all OXPHOS. Six replicates for each condition in any given experiment were used.

#### Cellular mitochondrial substrate oxidation measurement

Freshly isolated mice spleens were minced immediately, and functional mitochondria were isolated using the method previously reported (*39*). In brief, the minced cells were homogenized with a glass pestle in an isolation buffer (1M sucrose, 1M Tris–HCl, 1M KCl, 1M EDTA, and 10% BSA, pH 7.4). The homogenized tissue was centrifuged at 700*g* for 10 minutes at 4°C. The pellet was resuspended in buffer (1M sucrose, 0.1M EGTA-Tris, 1M Tris–HCl, pH 7.4) and centrifuged at 8000*g* for 10 minutes). Protein levels, measured by the BCA assay, were used to assess the mitochondrial concentration, and 2 μg of mitochondria were loaded into each well. ADP (A2754, Sigma), Oligomycin and FCCP, and Antimycin A (103015-100, Agilent Technologies, Santa Clara, CA) were subsequently injected, as described by the manufacturer. Cell Mito Stress Test was further combined with substrate pathway-specific inhibitors (etomoxir, UK5099, BPTES) (Agilent Technologies, Santa Clara, CA) to interrogate which of the three primary substrates (long chain fatty acids, glucose/pyruvate, and/or glutamine) fuel the mitochondria.

#### Disease blood monitoring

Approximately ~100μl of blood was collected via submandibular bleeding into EDTA-coated tubes. 1 ml of ACK buffer was used for erythrocyte lysis and then washed with PBS with 1% BSA and 2mM EDTA (FACS buffer). Cells were then stained with a cocktail of antibodies against CD45.2 (APC anti-mouse CD45.2 [104], Biolegend); CD45.1 (Brilliant violet 510™ anti-mouse CD45.1[A20], Biolegend); CD5(PE/Cy5 anti-mouse CD5 [53-7.3], Biolegend); B220 (Pacific Blue™ anti-mouse/human CD45R/B220 [RA3-6B2], Biolegend) CD3 (APC/Cy7 anti-mouse CD3 [17A2], Biolegend), CD11b (PE/Cy7 anti-mouse/human CD11b [M1/70], Biolegend) for 15 minutes at 4°C. Cells were further washed with FACS buffer and analyzed by flow cytometry. CLL-RT mice splenic B cells were further analyzed by CD71(Anti-CD71 Rat Monoclonal Antibody (PE (Phycoerythrin)) [clone: RI7217], Brilliant Violet 711™ anti-mouse CD21/CD35 (CR2/CR1) Antibody and Anti-CD38 Rat Monoclonal Antibody (PE (Phycoerythrin)/Cy7®) [clone: 90].

#### Immunohistochemistry staining

Freshly isolated spleens and bone marrow tissues were fixed in neutral formalin overnight and replaced with 70% ethanol the next day until the tissues were processed. Spleens were paraffin-embedded, and 10 μm sections were made for IHC staining. Ki67, CD5, B220 and cMYC levels were estimated by respective antibodies as reported (*37*) and horseradish peroxidase (HRP) conjugated secondary antibody to reveal the diaminobenzidine (DAB) staining. IHC stains were performed on Ventana Discovery Ultra (Ventana Medical Systems, Roche Diagnostics, Indianapolis, USA) IHC Auto Stainer. Briefly, the FFPE tissue blocks were sectioned at a thickness of 5 μm and put on positively charged glass slides. The slides were loaded on the machine, and deparaffinization, rehydration, endogenous peroxidase activity inhibition, and antigen retrieval were first performed. Then, each primary antibody was incubated, followed by DISCOVERY anti-Rabbit H.Q. or DISCOVERY anti-Mouse H.Q. and DISCOVERY anti-HQ-HRP incubation. The stains were visualized with DISCOVERY ChromoMap DAB Kit, counterstained with hematoxylin (Ventana), and coverslipped. IHC image analysis was done using Visiopharm tool. 4 areas were randomly chosen and quantified using in-house developed apps in the tool following the manufacturer guidelines.

#### Mouse serial transplantation

Mouse transplantation studies were conducted on 6-10-week-old immunocompetent CD45.1 C57BL/6 (sub-lethally irradiated with 400 cGy) or immunodeficient NSG mice using viably cryopreserved splenic cells from CLL animals. One million CLL cells were resuspended in 100-150μl of PBS and injected intravenously into CD45.1 C57BL/6 or NSG mice. For RT mice, 1 million cells were resuspended in 100-150μl of PBS and injected intravenously into NSG mice. CLL/RT burden in the peripheral blood was monitored by flow cytometry. The criteria for euthanasia were hunched back, lethargic movement, breathing difficulties, and leg paresis.

#### Transmission electron microscopy (TEM) of mouse and cultured cells

Cells were first fixed with 2.5% glutaraldehyde, 0.1M cacodylate buffer (Na (CH3)_2_AsO_2_·3H_2_O), pH7.2, at 4°C overnight. TEM sample preparation included post-fixation with osmium tetroxide, serial dehydration with ethanol, and embedment in eponate (*40*). Ultra-thin sections (70 nm thick) were acquired by ultramicrotomy and were examined on an FEI Tecnai 12 transmission electron microscope equipped with a Gatan OneView CMOS camera.

#### Cell proliferation measurement

Cells were cultured at a density of 5000 cells/100μl in RPMI media with 10% FBS in a 96 well-tissue culture plate. Cell proliferation was measured by CCK8 assay (Dojindo) for 4 days, measured every days. CCK8 absorbance was measured at 450nm. All measurements were made in triplicates, and the data were normalized to Day1.

#### mtDNA analysis

mtDNA copy number and amount were determined using qPCR-based and Pico Green-based flow cytometry assays. Genomic DNA was first extracted from mouse spleen tissue or cell lines using M.N. tissue DNA kit (Macherey Nagel, Allentown, PA). Primers were designed for mtDNA copy number measurement for mitochondrial gene mt-ND1 and nuclear genes *GAPDH* and *HMGB1*. 10ng of genomic DNA was used for qPCR, performed in replicates, and all data were normalized to nuclear DNA. The 2^^(-Delct)^ method determined the quantification of mtDNA.

#### Western Blotting

Cells were lysed in RIPA buffer (ThermoFisher Scientific, Waltham, MA) supplemented with a protease-phosphatase cocktail (Pierce™ Protease and phosphatase inhibitor minitablets EDTA-free, ThermoFisher Scientific, Waltham, MA) for 30 minutes at 4°C before sonication and protein quantification was measured using BCA assay (Pierce™, ThermoFisher Scientific, Waltham, MA). 20 μg protein was separated on SDS-PAGE (4-15% Criterion Precast Gel, Bio-Rad Laboratories, Hercules, CA) and transferred to nitrocellulose membrane (Trans-Blot Turbo nitrocellulose membranes, Bio-Rad Laboratories, Hercules, CA). Membrane strips were blocked in 5% BSA in TBS-0.1% Tween 20 and incubated overnight at 4°C with respective primary antibodies. Then, membranes were washed three times with TBS-0.1% Tween 20 and incubated for 1 hour with anti-mouse/rabbit secondary antibodies (ThermoFisher Scientific, Waltham, MA). Subsequently, the membranes were developed for ECL detection (Clarity Western ECL substrate, Bio-Rad Laboratories, Hercules, CA) following the manufacturer’s instructions. Images were acquired using ChemiDoc MP (Bio-Rad Laboratories, Hercules, CA). Protein bands were quantified using Bio-Rad imaging software (Image Lab 6.1).

#### Cytosolic and nuclear extraction

Cells were subjected to lysis using NE PER Nuclear and Cytoplasmic extraction kit (Thermo Fisher Scientific Inc, Grand Island, NY) and proteins were extracted following manufacturer guidelines.

#### Mitochondrial protein isolation

Cells were subjected to lysis using Pierce™ Mitochondria isolation kit for cultured cells (Thermo Fisher Scientific Inc, Grand Island, NY) and mitochondrial proteins were extracted following manufacturer guidelines.

#### *In vivo* luciferase experiments

For *in vivo* luciferase experiments, 5×10^5^ cells were first injected by tail-vein injection into recipient NSG mice. Bioluminescent imaging was performed at indicated time points by 50mg/kg intraperitoneal luciferin injection (Goldbio, St Louis, Missouri). Imaging was performed after 10 minutes using an IVIS imager (Spectral Instruments, Tucson, AZ). Bioluminescent radiance was quantified using Aura imaging software (Spectral Instruments, Tucson, AZ) with standard regions of interest in rectangles.

#### Drug administration *in vivo*

For drug administration experiments, 1 million murine RT cells were resuspended in 100-150μl of PBS and injected intravenously into NSG mice (vehicle (n = 13) and AZ5576 treated (n = 13), TTFA (n = 4) and AZ5576 + TTFA (n=5)). AZ5576 was dissolved in B-cyclodextrin solution and administered orally at a 60mg/kg dose twice 7 days apart. TTFA was dissolved in 2% DMSO and 30% PEG-300 solution. TTFA was administered orally at 25mg/kg 5 days a week. Same dosage was used for the combination. Treatment with either TTFA or AZ5576 or combination started 7 days post-RT engraftment. The criteria for euthanasia were hunched back, lethargic movement, breathing difficulties, and leg paresis.

#### RNA sequencing (RNA-seq), data processing, and differentially expressed mRNA analysis

Total RNA was isolated using Nucleospin RNA plus kit (Machery Nagel, Allentown, PA). Libraries for RNA-seq were constructed using the Stranded Total RNA Prep with Ribo-Zero Plus Kit (Illumina) and sequenced on Novaseq S4 platform using paired-end 150 bp mode. The fastq sequence files exported by sequencer were checked using FastQC (http://www.bioinformatics.bbsrc.ac.uk/projects/fastqc). Adaptors and low-quality bases were removed from the sequencing reads using Trimmomatic (*41*). The remaining reads were then aligned to the mouse reference genome (mm10) using STAR(*42*) with default parameters. Adaptor trimming and mapping quality reports were generated using MultiQC (*43*). The DESeq2(*44*) R package performed differential expression mRNAs analyses. mRNAs with absolute log_2_FC more than 1 and FDR less than 0.05 were identified as significantly dysregulated genes. For the semi-quantitative measurement of transcription, 2μg of total RNA was reverse transcribed using a high-capacity cDNA synthesis kit (Invitrogen Carlsbad, CA) with random hexamers following the manufacturer’s instructions. 1μl of cDNA was analyzed using Quant studio qPCR machine (Applied Biosystems, Bedford, MA), and transcript levels were quantified using the 2^^^(-ΔΔCt) method.

#### Gene expression cluster analysis between murine CLL and RT cells and human leukemia and lymphoma cell lines

Genes names of murine RNA-seq data were first transferred to human orthologues according to Ensembl 107 release, and then the RNA-seq counts mapped to each gene were combined with previously published human RNA-seq leukemia and lymphoma panel (*45*) to evaluate their similarity with other types of B-cell lymphomas. The RNA-seq counts were first normalized with DEseq2 followed by batch effect correction with combat. Genes in mitochondrial and sex chromosomes and those with maximum count lower than 5 were further removed. Then the top 1000 genes with highest variance were selected followed by removal of those genes with median absolute deviation (MAD) larger than 0.75. The data for the remaining 370 genes were used to generate heatmap plot using pheatmap package with Ward.D2 method and to calculate relationship using Pearson correlation with hierarchical clustering method in Rstudio.

#### Statistical analysis

Statistical analysis was performed using GraphPad Prism 9.3.1 (San Diego, California, USA). p values were calculated with one-way or two-way ANOVA test followed by Dunnett, Tukey, and Sidak multiple comparison test. A p-value < 0.05 was considered statistically significant. The type of statistical test used and the results, including p-value, means, median, and standard error, are shown in the figures and figure legends.

## CONTACT FOR REAGENT AND RESOURCE SHARING

Further information and requests for resources and reagents should be directed to and will be fulfilled by the Lead Contact, Lili Wang.

