## Supplementary files for "*MGA* deletion leads to Richter’s transformation via modulation of mitochondrial OXPHOS"

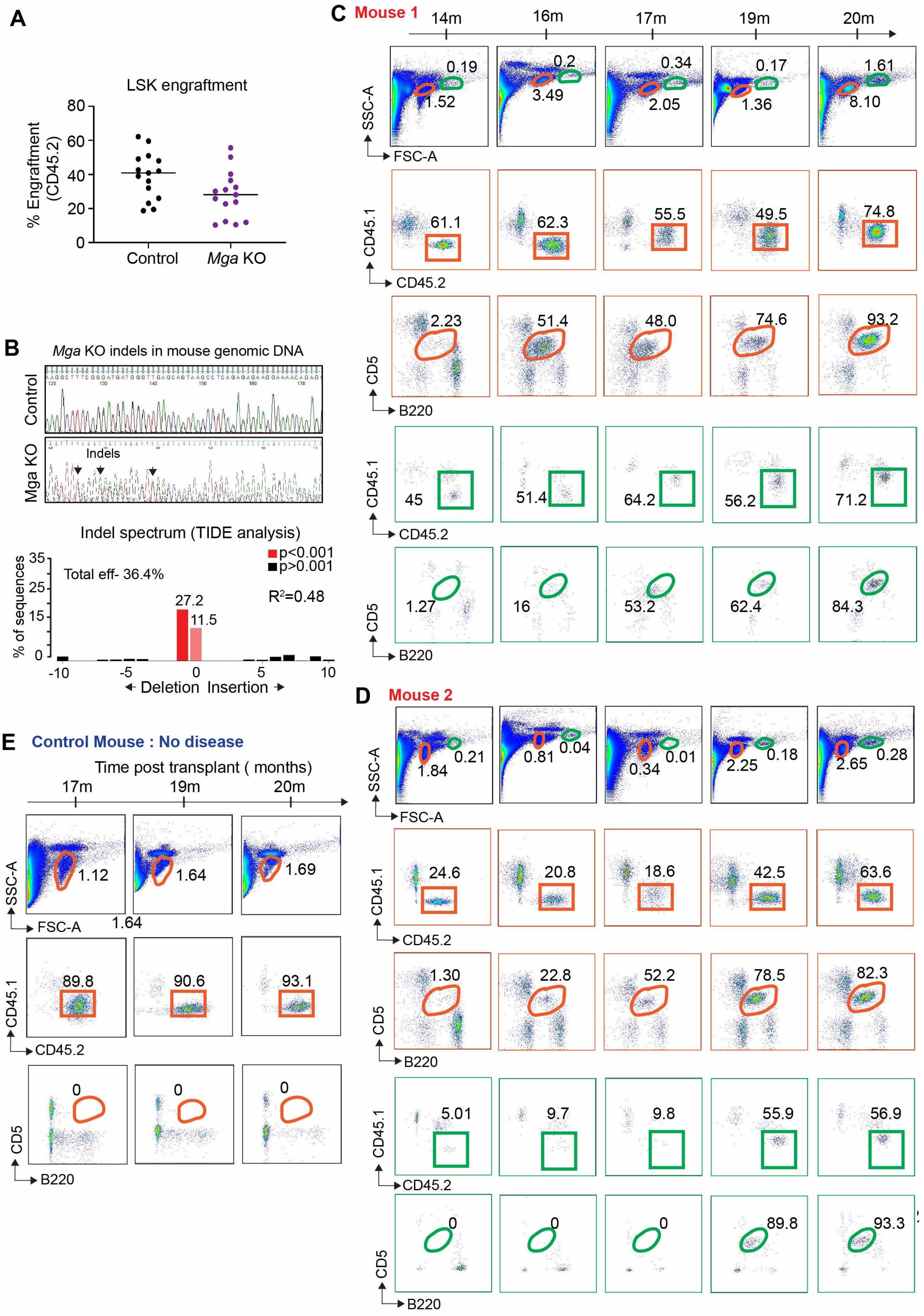
A
LSK engraftment
% Engraftment (CD45.2)
0
20
40
60
80
Control
Mga KO
B
Mga KO indels in mouse genomic DNA
Control
Mga KO
Indels
Indel spectrum (TIDE analysis)
Total eff- 36.4%
p<0.001
p>0.001
R²=0.48
27.2
11.5
% of sequences
35
25
15
0
-10
-5
0
5
10
Deletion Insertion
C
Mouse 1
14m
16m
17m
19m
20m
SSC-A
FSC-A
0.19
1.52
0.2
3.49
0.34
2.05
0.17
1.36
1.61
8.10
CD45.1
CD45.2
61.1
62.3
55.5
49.5
74.8
CD5
B220
2.23
51.4
48.0
74.6
93.2
45
51.4
64.2
56.2
71.2
1.27
16
53.2
62.4
84.3
D
Mouse 2
1.84
0.21
0.81
0.04
0.34
0.01
2.25
0.18
2.65
0.28
24.6
20.8
18.6
42.5
63.6
1.30
22.8
52.2
78.5
82.3
5.01
9.7
9.8
55.9
56.9
0
0
0
89.8
93.3
E
Control Mouse : No disease
Time post transplant ( months)
17m
19m
20m
1.12
1.64
1.69
1.64
89.8
90.6
93.1
0
0
0

**Figure S1. Validation of *Mga* KO in CLL-to-RT model. A,** Graph showing the percentage of engrafted donor CD45.2 LSK cells transduced with sgRNA targeting to control or *Mga* 9 months post engraftment. Each dot represents a mouse for the respective groups, and Y-axis shows the percentage of CD45.2 donor cells 9 months post engraftment. **B**, Sanger sequencing electropherogram showing the position of indels generated by *Mga* deletion in murine genomic DNA. Below is the % efficiency of *Mga* deletion by TIDE (Tandem decomposition of indels) analysis. **C-D**, Representative mouse showing the engraftment rate of genetically manipulated donor (CD45.2) cells and accumulation of $B220^{+}CD5^{+}$ CLL cells monitored by flow cytometry analysis of peripheral blood in the recipient mice (CD45.1). Green box indicates the CD45.2 *Mga* KO donor cells in the large lymphocytes, while orange box indicates the CD45.2 *Mga* KO donor cells in the small lymphocytes. Green circle indicates the $B220^{+}CD5^{+}$ in the large-sized lymphocytes, while orange circle indicates the $B220^{+}CD5^{+}$ in the small-sized lymphocytes. **E,** Representative mouse showing the engraftment rate (orange box) of a genetically edited mouse with control sgRNA and the accumulation of $B220^{+}CD5^{+}$ cells in the peripheral blood of recipient mice (orange circle).

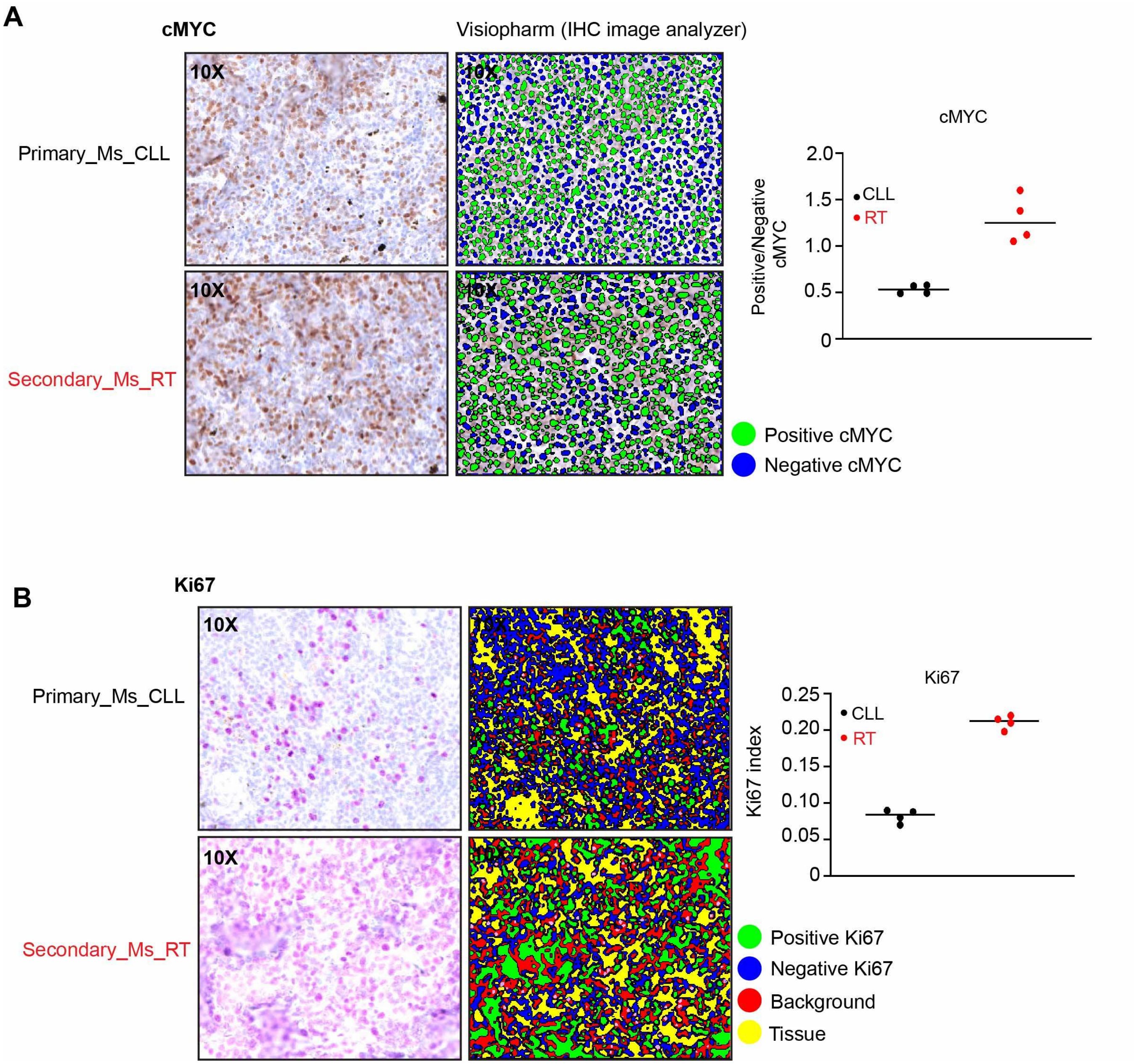


**Figure S2. Quantification of MYC and Ki67 proliferation index in murine spleen section derived from CLL and RT mice. A-B,** Representative image showing the MYC (A) and Ki67 (B) immunohistochemical staining in the spleen section from murine CLL and RT (10X magnification). AI (Artificial intelligence) based tool-Visiopharm is used for quantification in 4 randomly chosen areas. Graph indicates the ratio of positively stained MYC or Ki67 to negatively stained cells on the spleen section from mouse with CLL or RT.

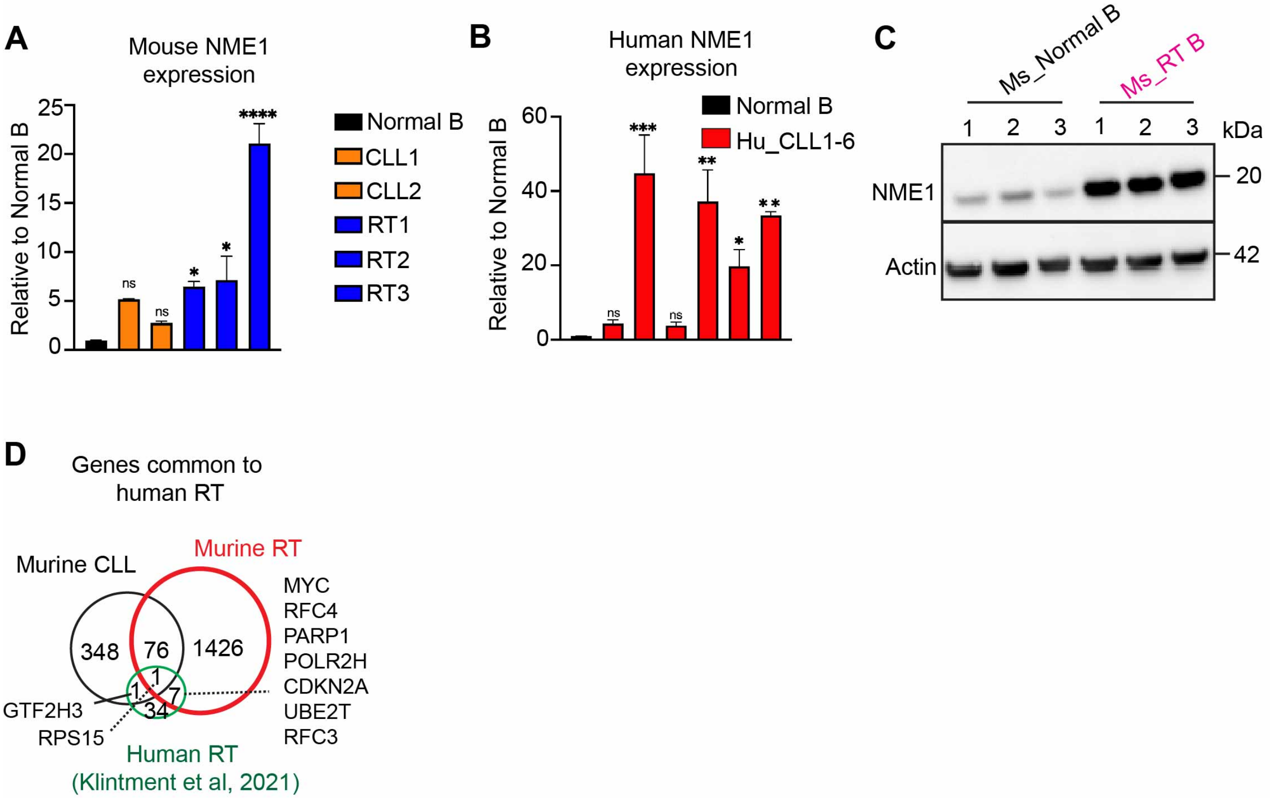


**Figure S3. Gene expression analysis from murine and human RT cells. A,** qPCR-RT validation of *NME1* expression in murine CLL, RT with respect to no disease mouse B cells. Y-axis represents the fold change expression with respect to murine normal B cells. **B,** qPCR-RT validation of *NME1* expression in human CLL with respect to normal B cells derived from healthy donors. Y-axis represents the fold change expression with respect to human normal B cells. **C,** NME1 expression in splenic B cells derived from three RT mice along with normal mouse B cells detected by immunoblotting. Actin is used as a loading control. **D,** Venn diagram showing the overlap of *Mga* deleted murine CLL- and RT- upregulated genes**,** and human RT upregulated genes with respect to nodal CLL (Klintment et al, 2021). $^{****}P< 0.0001$, $^{***}P<0.001$, $^{**}P<0.01$, $^{*}P<0.05$.

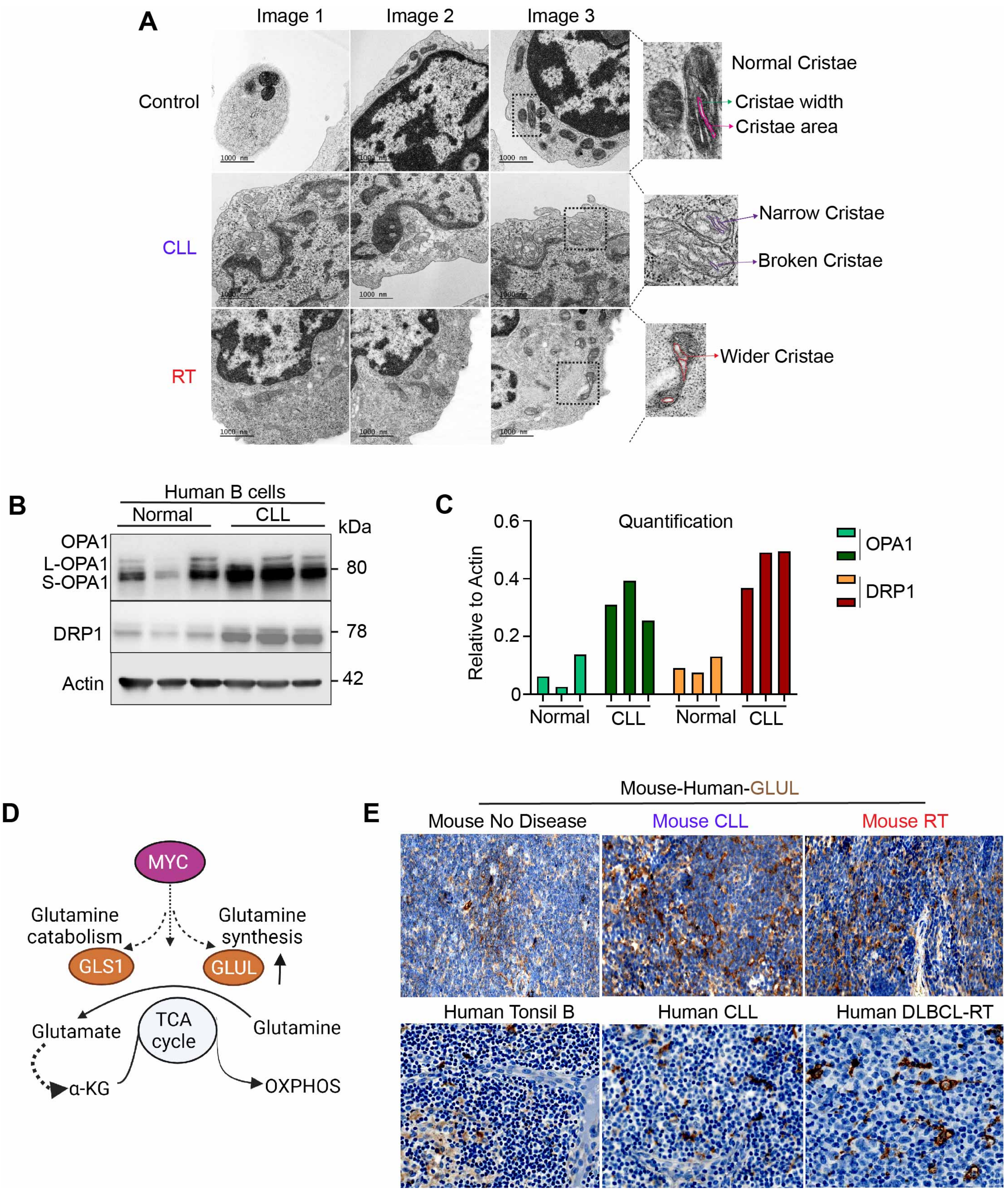

A
Image 1
Image 2
Image 3
Control
CLL
RT
Normal Cristae
Cristae width
Cristae area
Narrow Cristae
Broken Cristae
Wider Cristae
B
Human B cells
Normal
CLL
kDa
OPA1
L-OPA1
S-OPA1
80
DRP1
78
Actin
42
C
Quantification
Relative to Actin
0.6
0.4
0.2
0
Normal
CLL
Normal
CLL
OPA1
DRP1
D
MYC
Glutamine catabolism
Glutamine synthesis
GLS1
GLUL
TCA cycle
Glutamate
Glutamine
α-KG
OXPHOS
E
Mouse-Human-GLUL
Mouse No Disease
Mouse CLL
Mouse RT
Human Tonsil B
Human CLL
Human DLBCL-RT

**Figure S4. Mitochondrial dysregulation in murine and human RT cells. A,** Additional representative electron micrographs (EM) of mouse splenic B cells from control (no disease), CLL, and RT mice. (Scale bar: 1000 nm). **B,** Proteins involved in mitochondrial function (OPA1, DRP1) detected by immunoblotting in normal and CLL B cells derived from healthy donors and CLL patients, respectively. **C**, Quantification of OPA1 and DRP1 protein expression from B relative to Actin using ImageJ. **D,** Schema showing MYC regulating glutamine metabolism genes - GLUL (glutamine synthesis) and GLS1 (glutamine breakdown). **E,** Immunohistological staining of GLUL in mouse and human tissues. Top: splenic sections derived from mice with no disease, CLL, or RT. Bottom: tonsil from healthy donor and lymph node biopsies from two CLL-RT cases. 40x magnification with scale bar as 100μm.

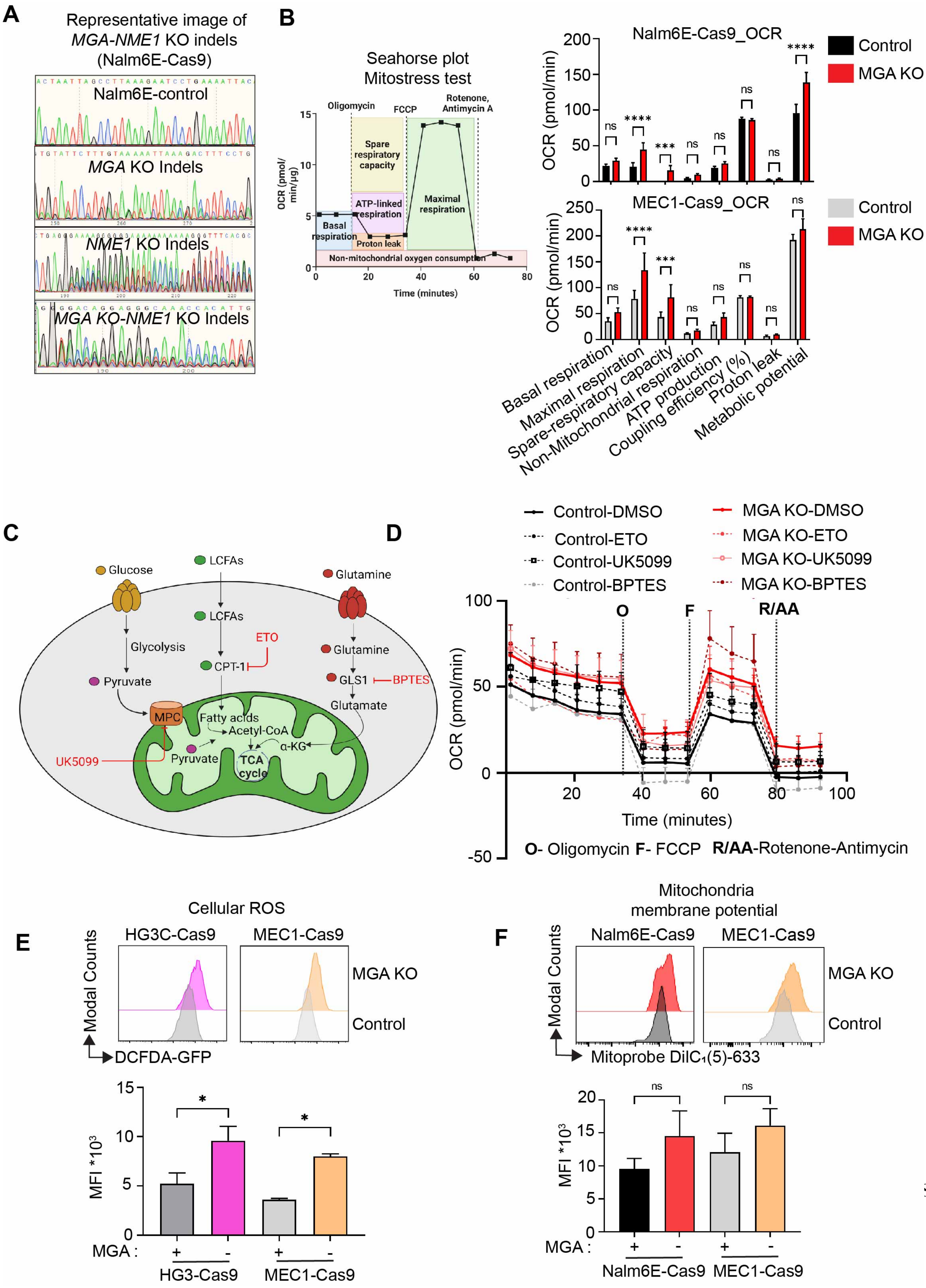
A
Representative image of
MGA-NME1 KO indels
(Nalm6E-Cas9)
Nalm6E-control
MGA KO Indels
NME1 KO Indels
MGA KO-NME1 KO Indels
B
Seahorse plot
Mitostress test
Oligomycin
FCCP
Rotenone, Antimycin A
Spare respiratory capacity
ATP-linked respiration
Maximal respiration
Basal respiration
Proton leak
Non-mitochondrial oxygen consumption
OCR (pmol/min/µg)
Time (minutes)
Nalm6E-Cas9_OCR
MEC1-Cas9_OCR
Control
MGA KO
OCR (pmol/min)
Basal respiration
Maximal respiration
Spare-respiratory capacity
Non-Mitochondrial respiration
ATP production
Coupling efficiency (%)
Proton leak
Metabolic potential
C
Glucose
LCFAs
Glutamine
Glycolysis
Pyruvate
CPT-1
ETO
GLS1
BPTES
Glutamate
MPC
UK5099
Fatty acids
Acetyl-CoA
α-KG
TCA cycle
D
Control-DMSO
Control-ETO
Control-UK5099
Control-BPTES
MGA KO-DMSO
MGA KO-ETO
MGA KO-UK5099
MGA KO-BPTES
O
F
R/AA
OCR (pmol/min)
Time (minutes)
O- Oligomycin F- FCCP R/AA-Rotenone-Antimycin
E
Cellular ROS
HG3C-Cas9
MEC1-Cas9
MGA KO
Control
Modal Counts
DCFDA-GFP
MFI *10³
MGA :
HG3-Cas9
MEC1-Cas9
F
Mitochondria
membrane potential
Nalm6E-Cas9
MEC1-Cas9
MGA KO
Control
Modal Counts
Mitoprobe DilC₁(5)-633
MFI *10³
MGA :
Nalm6E-Cas9
MEC1-Cas9

**Figure S5. Mitochondrial dysregulation in Nalm6E cells and MEC1 with and without *MGA*. A,** Representative sanger sequencing electropherogram of *MGA* KO, *NME1* KO, and *MGA*KO *NME1* KO indels in Nalm6E-Cas9 cells. **B,** Schema of inhibitors used in the substrate oxidation test by Seahorse Mito fuel flex analysis. Metabolic parameters measured by Seahorse Mitostress assay in MEC1-Cas9 (top) and Nalm6E-Cas9 (bottom) cells with or without *MGA*. **C-D,** Metabolic dysregulation in Nalm6E-Cas9 cells with *MGA* KO measured by the Seahorse Mitostress assay. Oxygen consumption rate (OCR) measured by Seahorse Mito Fuel Flex analysis from Nalm6E control, *MGA* KO, in the presence or absence of fuel pathway inhibitors. Etomoxir (ETO): long-chain fatty acid inhibitor; BPTES: glutamine inhibitor; UK5099: glucose inhibitor. **E,** Cellular ROS measured by DCFDA flow cytometry-based assay in HG3 and MEC1 cells with or without *MGA*. **F,** Mitochondrial membrane potential was measured by flow cytometry using membrane-potential-sensitive dye DilC1(5) coupled with disrupter CCCP. D $^{****}P<$ 0.0001, $^{***}P<0.001$, $^{**}P<0.01$, $^{*}P<0.05$.

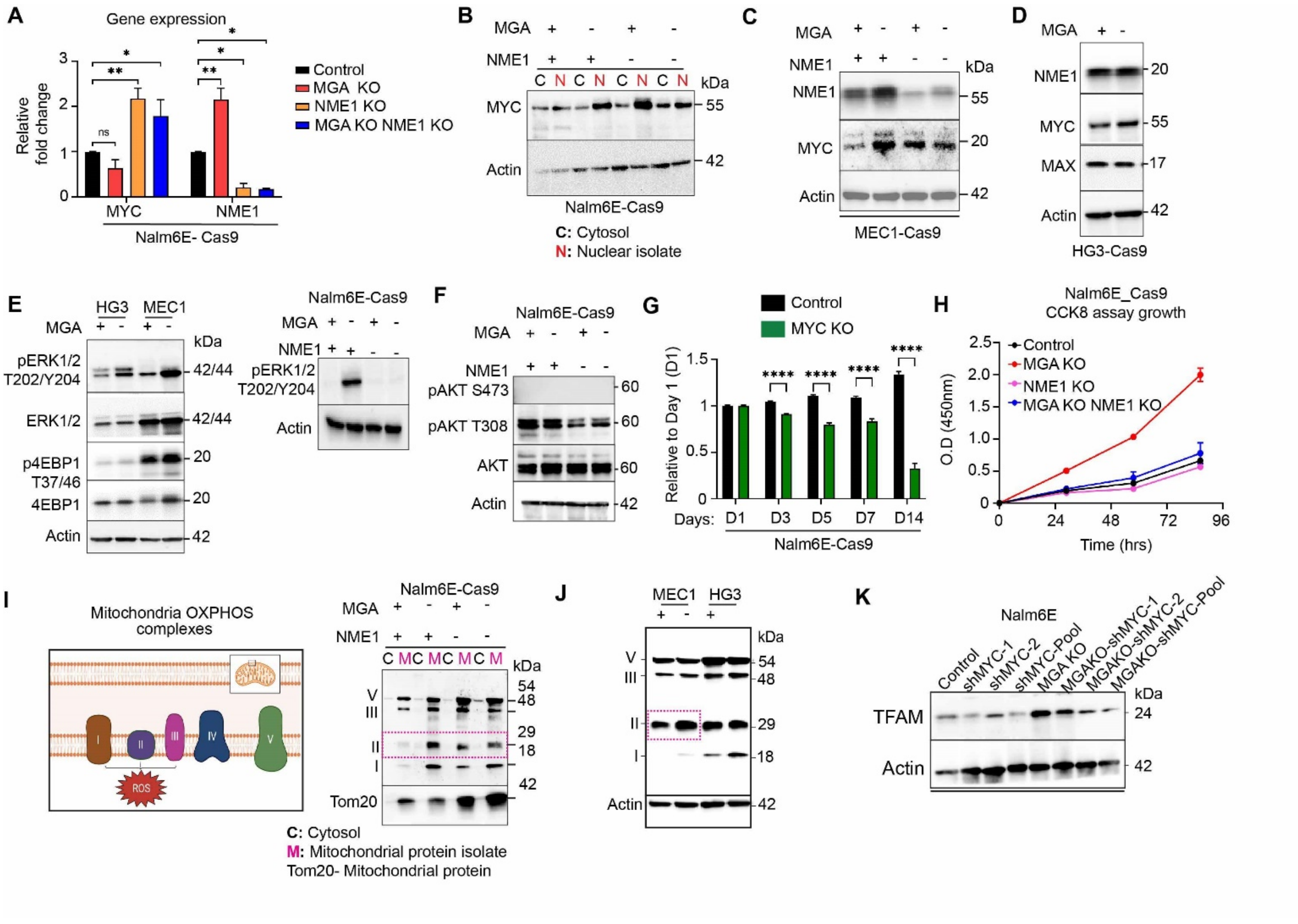


**Figure S6.** ***MGA* KO results in the activation of ERK and mTOR pathways in various cell lines.** A, qPCR-RT validation of MYC and NME1 in Nalm6E cells with different genotypes. Y axis indicates the fold change in gene expression with respect to control. **B,** MYC protein expression in the cytosolic and nuclear fractions of Nalm6E-Cas9 cells detected by immunoblotting. Actin is used as a control for the cytosolic fraction. **C,** NME1, and MYC protein expression were detected by immunoblotting in MEC1 cells. **D,** MYC, MAX, and NME1 protein expression was detected by immunoblotting in HG3-Cas9 cells with and without *MGA*. **E,** Phospho-ERK1/2 (T202/Y204), ERK1/2, phospho-4E-BP1 (T37/46), and 4E-BP1 protein expression in HG3 and MEC1 Cas9 cell lines with and without *MGA*. Phospho-ERK1/2 (T202/Y204) expression in Nalm6E-Cas9 cells with different genotypes. **F,** pAKT S473, and T308, total AKT protein expression detected by immunoblotting in Nalm6E-Cas9 cells with different genotypes. **G,**

Growth competition assay was performed using control Nalm6E-Cas9 cells and MYC KO cells. Cell growth was monitored by flow cytometry over 14 days. Percentage of cells was plotted. ****P< 0.0001, ns, Not significant. **H,** Cell growth was analyzed by CCK8 assay in Nalm6E cells with different genotypes over four days. Y axis indicates absorbance at 450nm. **I,** Cartoon showing the structure of mitochondrial OXPHOS complex proteins. Mitochondrial ETC OXPHOS complex protein expression in cytosolic and mitochondrial fractions of Nalm6E-Cas9 with different genotypes detected by immunoblotting. Tom20 is the outer mitochondrial protein used as a loading control. Increased complex II in *MGA* KO cells is highlighted in a pink box (dotted). **J,** Mitochondrial ETC (Electron transport chain) protein expression in MEC1 and HG3 cells with or without *MGA*. Increased complex II in MEC1 *MGA* KO cells is highlighted in a pink box (dotted). **K**, TFAM (transcription factor A, mitochondrial) protein expression in Nalm6E control with MYC knockdown (Control, shMYC1, shMYC2, pooled shMYC (1+2), and *MGA* KO, *MGA* KO shMYC1, shMYC2 and pooled shMYC (1+2).

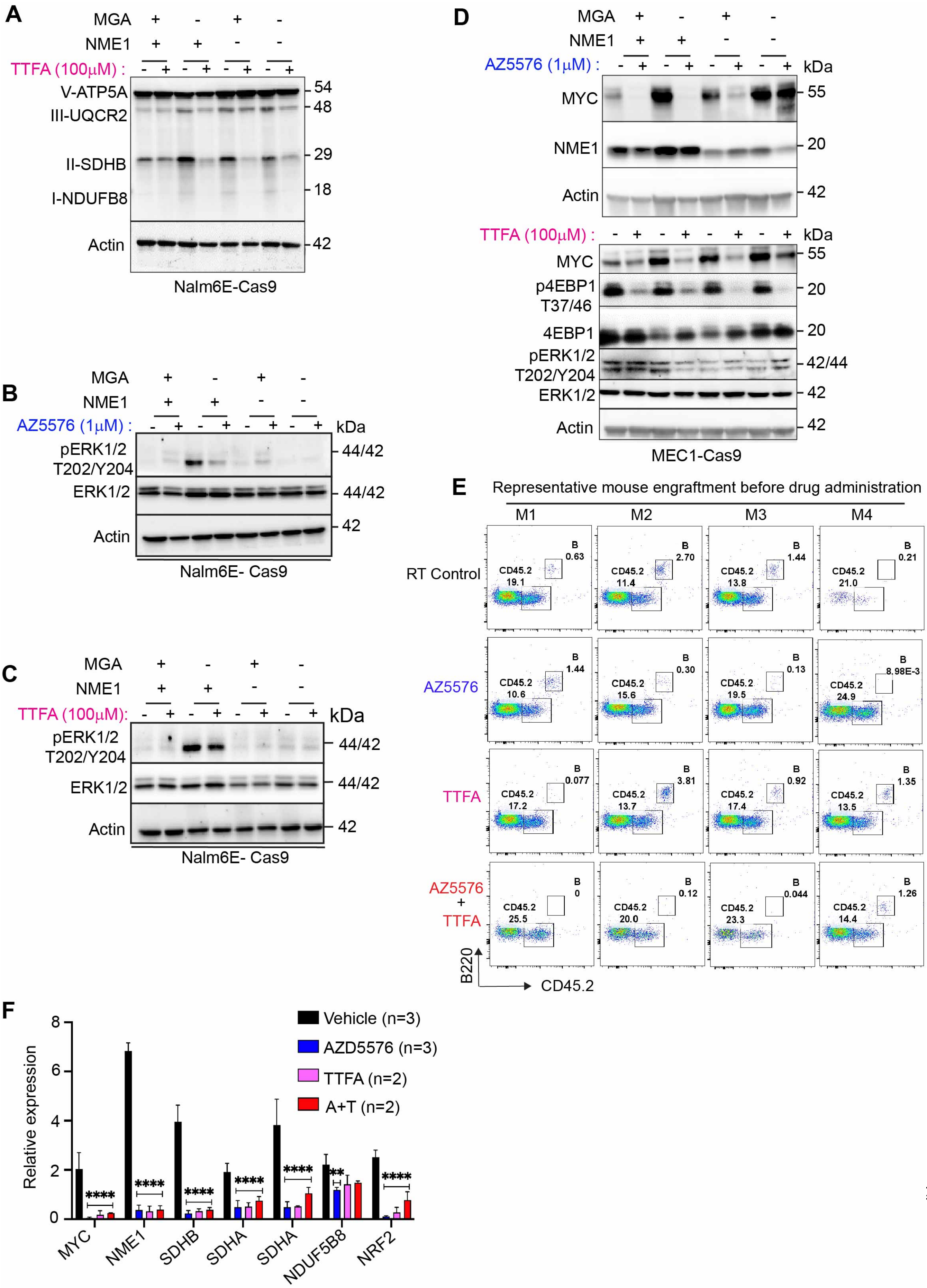
A
MGA + - + -
NME1 + + - -
TTFA (100μM) : - + - + - + - +
V-ATP5A
III-UQCR2
II-SDHB
I-NDUFB8
Actin
54
48
29
18
42
Nalm6E-Cas9
B
MGA + - + -
NME1 + + - -
AZ5576 (1μM) : - + - + - + - +
kDa
pERK1/2 T202/Y204
ERK1/2
Actin
44/42
44/42
42
Nalm6E- Cas9
C
MGA + - + -
NME1 + + - -
TTFA (100μM): - + - + - + - +
kDa
pERK1/2 T202/Y204
ERK1/2
Actin
44/42
44/42
42
Nalm6E- Cas9
D
MGA + - + -
NME1 + + - -
AZ5576 (1μM) : - + - + - + - +
kDa
MYC
NME1
Actin
55
20
42
TTFA (100μM) : - + - + - + - +
kDa
MYC
p4EBP1 T37/46
4EBP1
pERK1/2 T202/Y204
ERK1/2
Actin
55
20
20
42/44
42
42
MEC1-Cas9
E
Representative mouse engraftment before drug administration
M1 M2 M3 M4
RT Control
B 0.63 CD45.2 19.1
B 2.70 CD45.2 11.4
B 1.44 CD45.2 13.8
B 0.21 CD45.2 21.0
AZ5576
B 1.44 CD45.2 10.6
B 0.30 CD45.2 15.6
B 0.13 CD45.2 19.5
B 8.98E-3 CD45.2 24.9
TTFA
B 0.077 CD45.2 17.2
B 3.81 CD45.2 13.7
B 0.92 CD45.2 17.4
B 1.35 CD45.2 13.5
AZ5576 + TTFA
B 0 CD45.2 25.5
B 0.12 CD45.2 20.0
B 0.044 CD45.2 23.3
B 1.26 CD45.2 14.4
B220
CD45.2
F
Relative expression
0
2
4
6
8
Vehicle (n=3)
AZD5576 (n=3)
TTFA (n=2)
A+T (n=2)
****
****
****
****
****
**
****
MYC
NME1
SDHB
SDHA
SDHA
NDUF5B8
NRF2

**Figure S7. TTFA and AZD5576 treatment in cell lines and RT mice. A-D,** ETC complex, pERK, p4E-BP1 (mTORC1) pathway protein expression in Nalm6E and MEC1 cell lines with different genetic lesions upon treatment with TTFA, AZD5576 alone. **E,** RT cells were engrafted into NSG mice for one week, and engraftment was confirmed by flow cytometry of peripheral blood cells. **F,** Genes related to ETC complex and oxidative stress were tested in RT mice with TTFA and AZD5576 treatment by qPCR. $^{****}P<$ 0.0001, $^{**}P<0.01$.
